# ZNF512B binds RBBP4 via a variant NuRD interaction motif and aggregates chromatin in a NuRD complex-independent manner

**DOI:** 10.1101/2023.07.31.551264

**Authors:** Tim Marius Wunderlich, Chandrika Deshpande, Lena W. Paasche, Tobias Friedrich, Felix Diegmüller, Nadine Daus, Haniya Naseer, Sophie E. Stebel, Jörg Leers, Jie Lan, Van Tuan Trinh, Olalla Vázquez, Falk Butter, Marek Bartkuhn, Joel P. Mackay, Sandra B. Hake

**Author notes:** shared authors (Chandrika Deshpande, Lena W. Paasche, Tobias Friedrich). Institute of Biochemistry, Justus-Liebig University Giessen, 35392 Giessen, Germany (Sophie E. Stebel). Corresponding author: Sandra B. Hake, Institute for Genetics, Justus-Liebig-University Giessen, Heinrich-Buff-Ring 58-62, 35392 Giessen, Germany.

## Abstract

The evolutionarily conserved histone variant H2A.Z plays a crucial role in various DNA-based processes but the underlying mechanisms by which it acts are not completely understood.

Recently, we identified the zinc finger protein ZNF512B as an H2A.Z-, HMG20A- and PWWP2A-associated protein. Here, we report that ZNF512B binds the nucleosome remodeling and deacetylase (NuRD) complex. We discover a conserved amino acid sequence within ZNF512B that resembles the NuRD-interaction motif (NIM) previously identified in FOG-1 and other transcriptional regulators. By solving the crystal structure of this motif bound to the NuRD component RBBP4 and by applying several biochemical assays we demonstrate that this internal NIM is both necessary and sufficient for robust NuRD binding. Transcriptome analyses and reporter assays identify ZNF512B as a repressor of gene expression that can act in both NuRD-dependent and -independent ways. Surprisingly, high levels of ZNF512B expression lead to nuclear protein and chromatin aggregation foci that form independent of the interaction with the NuRD complex but depend on the zinc finger domains of ZNF512B. Our study has implications for diseases in which ZNF512B expression is deregulated, such as cancer and neurodegenerative diseases, and hint at the existence of more proteins as potential NuRD interactors.

## Introduction

Zinc finger (ZF) proteins are the most abundant protein family in eukaryotes and are characterized by the coordination of one or more zinc ions by a combination of cysteine and histidine residues (1,2). ZFs can bind a variety of molecules, such as DNA, RNA, lipids and proteins (3-6), and are important for many DNA-based processes involved in, for example, development, differentiation, metabolism signal transduction and immune responses (1). Numerous ZF proteins have been shown to act as transcription factors (TFs), often by facilitating the recruitment of other, enzymatically active proteins to specific DNA sequences. As a result, the structure of chromatin (7), which is the regulatory packaged form of the genetic material in eukaryotes, can be modified and functionally affected. The smallest unit of chromatin is the nucleosome: an octameric complex consisting of two of each of the four core histones H2A, H2B, H3 and H4 wrapped by approximately 150 bp of DNA (8). Different interconnected mechanisms have evolved to ensure the local and/or global control of nucleosome positions and DNA accessibility including the addition of chemical groups to DNA or histone proteins, ATP-dependent remodelling and incorporation of specialized histone variants (9).

One such intensively studied histone variant is the conserved and in higher eukaryotes essential histone variant H2A.Z (10). Its deposition into genomic regulatory regions, such as promoters and enhancers, leads to the control of transcription, genome stability and cell cycle progression, and plays an important role during stem cell differentiation and early development (11-13). It is thought that H2A.Z influences these processes by i) the deposition of the non-redundant paralogues H2A.Z.1 or H2A.Z.2.1 (or H2A.Z.2.2 in primates only) (14-16), ii) distinct post-translational modification patterns (12), iii) changes in nucleosome stability that depend on the presence or absence of other histone variants, such as H3.3 (17), and iv) the binding and recruitment of different chromatin-modifying complexes (18).

In previous studies, we have identified proteins enriched on H2A.Z-containing nucleosomes compared to those harbouring the replication-dependent histone H2A (16,19). Besides the known H2A.Z-specific chaperone complex SRCAP and the bromodomain protein BRD2, we found several chromatin-modifying complexes and functionally understudied proteins, such as PWWP2A, HMG20A and ZNF512B. We could show that PWWP2A is a direct H2A.Z nucleosome interactor required for head development. Furthermore, it binds to an MTA1-specific core nucleosome remodelling and deacetylase (NuRD) complex that we termed M1HR (20,21). This M1HR sub-complex lacks its remodelling CHD/MBD/GATA module due to competition between PWWP2A and MBD proteins for binding to MTA1 (22). In addition to M1HR, the PHD Finger 14 (PHF14), Retinoic Acid Induced 1 (RAI1), Transcription Factor 20 (TCF20) and High Mobility Box 20A (HMG20A) proteins have been repeatedly identified in both H2A.Z.1/2.1 and PWWP2A mass-spectrometry-based interactomes (16,19,20). In a recent follow-up study, we discovered that HMG20A is enriched at regulatory regions and controls important early head and heart transcription programs (23). HMG20A, in contrast to H2A.Z and PWWP2A, was able to pull down all components of the BHC/CoREST and NuRD repressive complexes, the latter including both remodelling (CHD) and deacetylase (HDAC) subunits. Within this and other quantitative mass-spectrometry screens, we also identified ZNF512B to be associated with H2A.Z, PWWP2A and HMG20A. ZNF512B, previously named GAM/ZFp (24), is a vertebrate-specific ZF protein (25). Few studies on ZNF512B functions are available. These demonstrate its role in gene regulation and cell homeostasis (25), indicate a possible association with NuRD in a mass-spectrometry based screen (26), and suggest a putative, yet unclear, role in the neurodegenerative disease Amyotrophic Lateral Sclerosis (ALS) (27-29).

Here, we show that ZNF512B can cause severe chromatin compaction when overexpressed and demonstrate that it can directly bind the NuRD complex. We discovered a conserved amino acid sequence within ZNF512B that resembles the functional NuRD-interaction motif (NIM) first identified in the transcription factor FOG-1 (also known as FOG-1-like motif) (30). Using a lysine to alanine point mutant, we demonstrate that this putative FOG-1-like motif in ZNF512B is a functional protein-protein interaction domain, which is both necessary and sufficient for NuRD binding but not required for chromatin compaction. The crystal structure of this motif bound to the NuRD component RBBP4 reveals a binding mode that is overall conserved with other RBBP4 complexes, although distinct differences have also been observed. A mutagenesis screen of the ZNF512B NIM confirms the observed interactions and indicates the existence of a variant internal NuRD binding sequence. Comparison of NIMs across many characterized NuRD interacting proteins reveals the presence of two different consensus sequences that appear to depend mainly on the protein location of the motif: i) an N-terminal RRKQxxP and ii) an internally located RKxxxPxK/R motif.

Moreover, we demonstrate that ZNF512B acts as transcriptional repressor and that its overexpression as well as its endogenous depletion result in the deregulation of gene expression. We further show that elevated ZNF512B levels cause chromatin aggregation in a NuRD-independent but ZF-dependent manner, possibly due to its ability to bind DNA and to di- or oligomerize.

In summary, ZNF512B binds directly to the NuRD complex, regulates transcription and, when present in elevated levels, affects chromatin architecture.

## Materials and Methods

### Cell culture and transfections

HeLa Kyoto (HeLaK), U2OS, HEK293 and HEK293T cells were cultured in Dulbecco’s modified Eagle medium (DMEM, Gibco), HCT116 cells were cultured in McCoy’s 5A modified Medium, both supplemented with 10% fetal calf serum (FCS; Gibco) and 1% penicillin/streptomycin (37^°^C, 5% CO_2_) and routinely tested via PCR for mycoplasma contamination. Transfections of cells were performed using FuGENE® HD Transfection Reagent (Promega) according to the manufacturer’s instructions (Promega). 48 hours after transfection, cells were harvested by trypsinization for various assays. Sf9 cells were cultured in Sf-900TM II SFM medium (Gibco) and maintained at 27^°^C and 90 rpm.

### Cloning of ZNF512B and its mutants

To generate human ZNF512B (mutant) plasmids, total RNA of HeLaK cells was purified using RNeasy-Kit (QIAGEN) and reverse transcribed using Transcriptor First Strand cDNA Synthesis Kit (Roche). cDNA was amplified by Q5® High-Fidelity DNA Polymerase (New England Biolabs) and cloned into pIRESneo-GFP (31), pEGFP-N2, p3xFLAG-CMV-10, pFastBac1 (Invitrogen) and pAB-Gal94 (32) vectors. Point mutation constructs were generated via site-directed mutagenesis (33).

### Primers and Oligos

All primers and oligos are listed in the Supplementary Information.

### Immunofluorescence microscopy

Immunofluorescence staining and microscopy was performed as previously described (34). Briefly, 1×10^5^ adherent cells expressing GFP, GFP-ZNF512B or its deletions/mutants were seeded on coverslips in 24-well plates and cultured overnight. The next day, cells were washed two times with PBS and fixed for 15 min in PBS containing 1% formaldehyde. After washing, cells were permeabilized and blocked with PBS containing 0.1% Triton^TM^-X-100 (PBS-T) and 1% bovine serum albumin (BSA) for 20 min. Cells were incubated stepwise with primary and then secondary antibodies in PBS-T + 1% BSA for 30 min with three wash-steps using PBS-T in between. After three final PBS-T wash-steps, DNA was stained with 10 µg/ml Hoechst solution for 3 min. After washing with H_2_O, coverslips with cells were then mounted in Fluoromount-G® mounting medium (SouthernBiotech) on microscope slides. Images were acquired using an Axio Observer.Z1 inverted microscope (Carl Zeiss, Oberkochen, Germany) with an Axiocam 506 mono camera system. Image processing was performed with Zeiss Zen 3.1 software (blue.edition). Intensity profiles were created using ImageJ2 (Version 1.9.0/1.53t) software.

### In situ staining for β-galactosidase activity

1×10^5^ HeLaK cells expressing GFP or GFP-ZNF512B or treated with Doxorubicin (100 nM, 2 days, Fisher Scientific) were seeded on coverslips in 24-well plates. *In situ* staining was done as described (35). In brief, after washing with PBS and fixing with 3.7% formaldehyde, cells were incubated overnight at 37°C in freshly prepared staining buffer (PBS pH 6.0, 2 mM MgCl_2_, 5 mM K_3_Fe[CN]_6_, 5 mM K_4_Fe[CN]_6_, 1 mg/ml X-Gal). After washing with H_2_O, coverslips with cells were mounted and imaged as described for immunofluorescence microscopy.

### Antibodies

All antibodies are listed in the Supplementary Information.

Polyclonal antiserum against ZNF512B was generated by expression of recombinant human His-ZNF512B protein in insect cell followed by FPLC-based purification (see EMSAs) followed by immunization of rabbits by BJ-Diagnostik BioScience GmbH.

### DNase immunoprecipitation (DNase-IP) and peptide competition

Preparation of DNase-digested cell extracts and immunoprecipitation was done using GFP-Trap® Magnetic Particles M-270 (Chromotek) following the manufacturer’s protocol. In brief, cells were incubated for 30 min on ice with RIPA buffer (10 mM Tris/Cl pH 7.5, 150 mM NaCl, 0.5 mM EDTA, 0.1% SDS, 1% Triton^TM^ X-100, 1% deoxycholate) supplemented with DNaseI (75 Kunitz U/ml, Thermo Fisher Scientific) and MgCl_2_ (2.5 mM). After centrifugation at 17,000 g for 10 min at 4°C, lysate was transferred to a fresh tube and mixed with Dilution buffer (10 mM Tris/Cl pH 7.5, 150 mM NaCl, 0.5 mM EDTA). Lysate was then incubated with GFP-Trap® Magnetic Particles M-270 rotating for 1 hour at 4°C. For peptide competition, lysate was incubated with peptide solution rotating for 1 hour at 4°C before incubation with beads. After incubation, beads were washed three times with Wash buffer (10 mM Tris/Cl pH 7.5, 150 mM NaCl, 0.05% Nonidet^TM^ P-40 Substitute, 0.5 mM EDTA), and precipitated proteins were eluted by boiling beads for 5 min in 2x SDS-sample buffer. Eluates were compared to input material via semi-dry Western blotting. Details on applied antibodies are listed in Supplementary Information.

### Fluorescence Polarization (FP)

To determine the dissociation constant (*k*_D_), serial dilutions of RBBP4 protein (4.5 - 0 µM, 45 µL) in 50 mM Tris-Cl pH 7.5, 150 mM NaCl and 0.02% Triton X-100 were added to 5 µL of the fluorescently-tagged tracer peptide Ac-SALL4 (2-12)-FAM for a final volume of 50 µL with 1.0 nM tracer. The 384-well plate (GBO 781 900) was measured (Tecan Spark 20M) after 60 min and the *k*_D_ values were calculated by converting the mP values into their corresponding anisotropy (A) values following the equation:

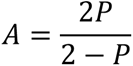

Herby, *A* = anisotropy and *P* = polarization. These values were plotted versus the respective concentrations of the protein with a non-linear regression according to the following equation (detailed instructions are found in (research.fredhutch.org/content/dam/stripe/hahn/methods/biochem/beacon_fluoresce nce_guide.pdf):

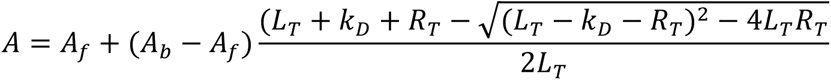

Hereby, *A* is the experimental anisotropy, *A_f_*is the anisotropy for the free ligand, *A_b_* is the anisotropy for the fully bound ligand, *L_T_* is the total added concentration of ligand, *k*_D_ is the equilibrium dissociation constant and *R_T_* is the total receptor concentration. For the competitive FP-based binding experiments RBBP4 protein was preincubated with tracer peptide in 50 mM Tris-Cl pH 7.5, 150 mM NaCl and 0.02% Triton X-100 for 60 min. To 5 µL of this protein-tracer complex, 5 µL of the corresponding peptide serial dilutions (1.5 – 0 mM) were added, with a final concentration of 1.0 nM tracer, 1.25 µM protein. After 60 min the polarization values were measured and plotted against the logarithmic peptide concentration to determine the inhibitory concentration (IC_50_) by fitting the dose-response to four parameter logistic (4-PL) curves in GraphPad Prism 6.

### MNase immunoprecipitation (MNase-IP) and MS sample preparation

Preparation of Micrococcal nuclease (MNase)-digested nuclear extracts was performed as previously described (16,19,20,34). Briefly, nuclei were isolated by incubation with PBS containing 0.3% Triton^TM^ X-100 for 10 min at 4°C and washed in PBS, before being resolved in 500 µl freshly prepared Ex100 buffer (10 mM HEPES pH 7.6, 100 mM NaCl, 1.5 mM MgCl_2_, 0.5 mM EGTA, 10% Glycerol, 10 mM ϕ3-Glycerol phosphate, 1 mM DTT, 2 mM CaCl_2_). Chromatin was then digested with 1.5 U MNase (sufficient for 4×10^7^ cells, Sigma-Aldrich) for 20 min at 26°C. Digestion was stopped by adding 10 mM EGTA and transferring the sample to 4°C. After centrifugation for 30 min at 13,000 rpm at 4°C, supernatant containing mostly mononucleosomes was transferred to a fresh reaction tube. To assess integrity of mononucleosomes, DNA was isolated from 25 µl of sample using the QIAquick PCR Purification Kit (QIAGEN) and subjected to agarose gel electrophoresis. For IP experiments, MNase-digested nuclear extracts were incubated overnight with GFP-Trap® Magnetic Particles M-270 (Chromotek) rotating at 4°C. Beads were washed twice with 1 ml Washing Buffer I (10 mM Tris pH 7.5, 150 mM NaCl, 0.1 % Nonidet^TM^ P-40 substitute and twice with 1 ml Washing Buffer II (10 mM Tris pH 7.5, 150 mM NaCl).

For immunoblot analysis, precipitated proteins were eluted by boiling the beads for 5 min in SDS-sample buffer. Eluates were compared to input material via semi-dry Western blotting. Details on applied antibodies are listed in Supplementary Information.

For label-free quantitative mass spectrometry (lf-qMS), precipitated proteins were eluted by incubation of beads with Elution Buffer (2 M Urea, 50 mM Tris pH 7.5, 2 mM DTT, 20 µg/ml Trypsin Gold (Promega)) for 30 min at 37°C shaking at 1,400 rpm in the dark. Eluted peptides in the supernatant were transferred into a fresh reaction tube. Peptides remaining bound to beads were alkylated/eluted in Alkylation Buffer (2 M Urea, 50 mM Tris pH 7.5, 10 mM chloroacetamide) for 5 min at 37°C shaking at 1,400 rpm in the dark. Both eluates were combined, and peptides were further alkylated and digested overnight at 25°C shaking at 800 rpm in the dark. Trypsin digestion was stopped by adding 1% trifluoroacetic acid (Thermo Fisher Scientific). Peptides were subjected to lf-qMS (see below). Lf-qMS experiments were performed twice (biological replicates) with four technical replicates each.

### Label-free quantitative mass spectrometry (lf-qMS)

#### MS data acquisition

After tryptic digest (see MNase IP), peptides were desalted on StageTips (36) and analysed by nanoflow liquid chromatography on an EASY-nLC 1000 system (Thermo Fisher Scientific) coupled online to a Q Exactive Plus mass spectrometer (Thermo Fisher Scientific). Peptides were separated on a C18-reversed phase capillary (25 cm long, 75 μm inner diameter, packed in-house with ReproSil-Pur C18-AQ 1.9 μm resin, Dr Maisch GmbH) directly mounted on the electrospray ion source. We used a 90 min gradient from 2% to 60% acetonitrile in 0.5% formic acid at a flow of 225 nl /min. The Q Exactive Plus was operated with a Top10 MS/MS spectra acquisition method per MS full scan.

#### MS data analysis

The raw files were processed with MaxQuant (37) (version 1.6.5.0) against a human uniprot database (81,194 entries). Carbamidomethylation was set as fixed modification while methionine oxidation and protein *N*-acetylation were considered as variable modifications. The search was performed with an initial mass tolerance of 7 ppm mass accuracy for the precursor ion and 20 ppm for the MS/MS spectra in the HCD fragmentation mode. Search results were processed with MaxQuant and filtered with a false discovery rate of 0.01. The match between run option and the LFQ quantitation were activated. Protein groups marked as reverse, contaminants, only identified by site and less than 2 peptides (at least 1 unique) were removed. Non-measured values were imputed with a beta distribution at the limit of quantitation. For creating the volcano plots, a p-value of a two-tailed Welch’s *t*-test was calculated. Plots were created using the ggplot2 package.

### Structure

#### Peptide Synthesis

A peptide comprising residues 419–430 of human ZNF512B (UniProt accession number Q96KM6) was chemically synthesized and purified by ChinaPeptides (Shanghai, China). The peptide was N-terminally acetylated and C-terminally amidated.

#### Expression and purification of recombinant RBBP4

Recombinant expression of full-length RBBP4 (residues 1–425; UniProt accession number Q09028) was carried out as described in (38). Briefly, the pFBDM vector carrying the full length RBBP4 with an N-terminal 6xHis tag and a thrombin protease cleavage site was used for expression in insect cells. Recombinant baculovirus generation and large-scale expression in Sf9 cells were carried out using the Bac-to-Bac™ (Invitrogen) expression system.

The harvested cells were resuspended on ice in lysis buffer (20 mM Tris pH 8.0, 150 mM NaCl,10 mM MgCl_2_, 1 mM β-mercaptoethanol, Complete® EDTA-free protease inhibitor tablet). Cells were lysed by sonication, NP-40 (0.1 % v/v) and DNase I (10 µg/mL) were added to the cell lysate and the lysate was then clarified by centrifugation. The supernatant was incubated with nickel-nitrilotriacetic acid beads (Qiagen) for 2 h at 4 °C. The beads were washed with lysis buffer containing increasing concentrations of imidazole (2, 20 and 40 mM) and RBBP4 was eluted with 500 mM imidazole. 6xHis-tag was cleaved by dialysing the eluted protein overnight at 4°C in 20 mM Tris (pH 8.0), 150 mM NaCl and 2.5 mM CaCl_2_, together with 500 U of thrombin. RBBP4 was further purified by size-exclusion chromatography, using a HiLoad™ 16/60 Superdex™ 75 pg column (Cytiva) in 20 mM Tris (pH 7.5), 150 mM NaCl and 1 mM DTT. Fractions containing purified RBBP4 were concentrated to 5 mg/ml and stored at -20°C until required.

#### Crystallization and Structure Determination

Synthetic ZNF512B peptide (residues 419-430) was dissolved in demineralized water at a concentration of 10 mM. Concentrated RBBP4 (5 mg/ml) was mixed with ZNF512B peptide at a molar ratio of 1:2 and incubated on ice for 2 h. Crystallization trials were set up at 4 °C in 96-well plates as sitting drops using 0.2 μl of protein solution and 0.1 μl crystallization solution. Single crystals were obtained in 0.1 M MES/imidazole, pH 6.5, 40% v/v ethylene glycol; 20 % w/v PEG 8000, 0.1 M mixture of carboxylic acids (Morpheus screen, Molecular Dimensions). The crystals were harvested into a cryoprotectant solution containing 25% glycerol in mother liquor before cryocooling in liquid nitrogen. X-ray diffraction data were collected on MX2 beamline (microfocus) at the Australian Synchrotron at 100 K and a wavelength of 0.9537 Å. The dataset was processed and scaled using XDS (PMID: 20124692) and Scala (PMID: 16369096) respectively, at 2.2 Å resolution. The RBBP4-ZNF512B structure was solved by molecular replacement using PHENIX phaser-MR (PMID: 12393927) with RBBP4-FOG-1 (PDB ID: 2xu7) as a search model without the FOG-1 or other ligands. The crystal belonged to space group P 1 21 1, with two molecules in the asymmetric unit. Further model building was performed by iterative rounds of manual model building in real space using COOT (39), followed by refinement using REFMAC5 (40) and PHENIX (41). Validation of the structure was carried out with MolProbity (42). Interface analysis was performed using the EBI PISA server (43) and PDBsum (44). Figures were generated using PyMOL (45).

### Electrophoretic Mobility Shift Assays (EMSAs)

Expression and purification of recombinant His-ZNF512B protein was done as described (34). Briefly, pFastBac1 vectors were transformed into DH10Bac bacteria and viruses were generated according to the Bac-to-Bac™ Baculovirus Expression System Protocol (Invitrogen). Extracts were prepared 3 days after infection by washing the cells twice with ice cold PBS and incubation on ice for 10 min in hypotonic buffer (10 mM HEPES-KOH pH 7.9, 1.5 mM MgCl_2_, 10 mM KCl). After brief vortexing and centrifugation, the pellet (nuclei) was incubated for 20 min in hypertonic buffer (20 mM HEPES-KOH pH 7.9, 25% glycerol, 420 mM NaCl, 1.5 mM MgCl_2_, 0.2 mM EDTA).

After centrifugation for 10 min at 16.000x g the supernatant was stored at -20^°^C or directly used for EMSAs. Alternatively, the supernatant was purified by FPLC using the Äkta Start (Cytiva) via a His-Trap column (71502768) by standard procedures outlined in the instructions of the manufacturer.

EMSAs were performed as originally described (46) with the exception that 300 ng of salmon sperm DNA per reaction were used as an unspecific competitor. Cy5 end-labelled unmethylated or Cy3 end-labelled methylated double-stranded oligos (in which all eight Cytosines in a CpG context were methylated) were used as EMSA probes; the sequence was derived from the mouse H19 promoter (for sequence details see Supplementary Information).

### Luciferase Reporter Assay

Transfections of HEK293T cells were performed using the CaPO4-method as previously described (47). Per well of a 6-well plate 1 µg of 4xUAS-tk-luc reporter plasmid, 0.5 µg expression vectors for expression of Gal-fusion proteins and 0.5 µg pCMV Luc vector for normalization were transfected. Cells were harvested 48 h after transfection, lysed and luciferase and β-Gal assays were conducted with an Orion L instrument (Berthold) were conducted.

### RNA interference (RNAi) mediated knock-down of ZNF512B

For RNAi-mediated knock-down, ON-TARGETplus SMARTpool siRNAs (Horizon Discovery, Cambridge, UK) targeting ZNF512B mRNA or a non-targeting control pool were transfected using DharmaFECT1 transfection reagent (Horizon Discovery, Cambridge, UK) according to the manufacturer’s instructions. Briefly, for a 6-well plate 1×10^5^ HelaK cells were seeded 24 h before transfection. 10 µl of 5 µM targeting or non-targeting siRNAs were mixed with 190 µl reduced serum medium (Opti-MEM™, Gibco). In a separate tube, 4 µl DharmaFECT1 were mixed with 196 µl Opti-MEM™. Both tubes were incubated for 5 minutes, before combining and incubating for further 20 minutes. In the meantime, cells were washed once with PBS followed by the addition of 1.6 ml DMEM supplemented with 10% FCS without antibiotics. The transfection reaction was added dropwise, and cells were cultured for two days before experiments were conducted.

### RNA extraction, RT-qPCR and RNA-seq

RNA extraction was performed as described (48). Briefly, total RNA was extracted using the RNeasy Mini Kit (QIAGEN) with on column DNase digest (RNase-Free DNase Set, QIAGEN) according to the manufacturer’s instructions. 1 µg total RNA was reverse transcribed using the Transcriptor First Strand cDNA Synthesis Kit (Roche) with random hexamer primers according to the manufacturer’s protocol. For subsequent qPCR analysis, technical triplicates of 6 µl cDNA (1:20 dilution), 7.5 µl iTaq™ Universal SYBR® Green Supermix (Bio-Rad) and 1.5 µl primer mix (5 µM forward and reverse primer) were used. The qPCR program consisted of 5 min at 95^°^C, followed by 40 cycles at 95^°^C for 3 s and 60^°^C for 20 s. Lastly, Ct values of technical triplicates were averaged, and fold change expression was calculated with the Delta-Delta Ct method, normalizing to GFP and HPRT1 expression.

Total RNA from different human tissues was commercially acquired from Applied Biosystems and BioChain and used as described previously (49).

The samples sizes for each experiment are given in the figures and legends. The methods of statistical analyses are also described in each figure legend.

### mRNA-seq Analysis

RNA sequencing (RNA-seq) was performed at Biomarker Technologies (Germany). Raw sequencing files (FASTQ) were adaptor and quality trimmed using trimGalore (50). Alignment of the trimmed sequencing reads to the hg19 reference genome (downloaded from Illumina’s iGenomes) was performed using Hisat2 v.2.2.1 with “-- min-intronlen 30 –max-intronlen 3000” parameters (51) and stored as binary alignment maps (BAM). Read count per gene tables based on BAM files were generated within R using the summarizeOverlaps function of the GenomicAlignments package (52). Normalization of read counts and detection of differentially expressed genes was calculated with DESeq2 (53) based on the read counts per gene tables. If not other indicated, significantly differentially expressed genes were chosen using a log2FC threshold of > 1 or < -1 and an adjusted p-value < 0.05. Identification of enriched biological pathway was performed using clusterProfiler (54). Volcano plots were generated with the EnhancedVolcano R package (55).

## Results

### ZNF512B overexpression leads to the formation of chromatin foci

Recently, we identified the interactomes of human H2A.Z nucleosomes, as well as of the H2A.Z-associated proteins PWWP2A and HMG20A using label-free quantitative mass spectrometry (lf-qMS) (16,19,20,23). One common protein identified in all screens was the zinc finger protein ZNF512B, which is ubiquitously expressed in all human tissues at low levels, with the highest mRNA content in brain and tumour samples (Supplementary Figure 1A). ZNF512B contains eight ZF domains, which are arranged in four pairs consisting of one atypical C2HC followed by one typical C2H2 ZF each, as well as a large internal region (I) that harbours a nuclear localization signal (NLS) (Figure 1A).

**Figure 1:**
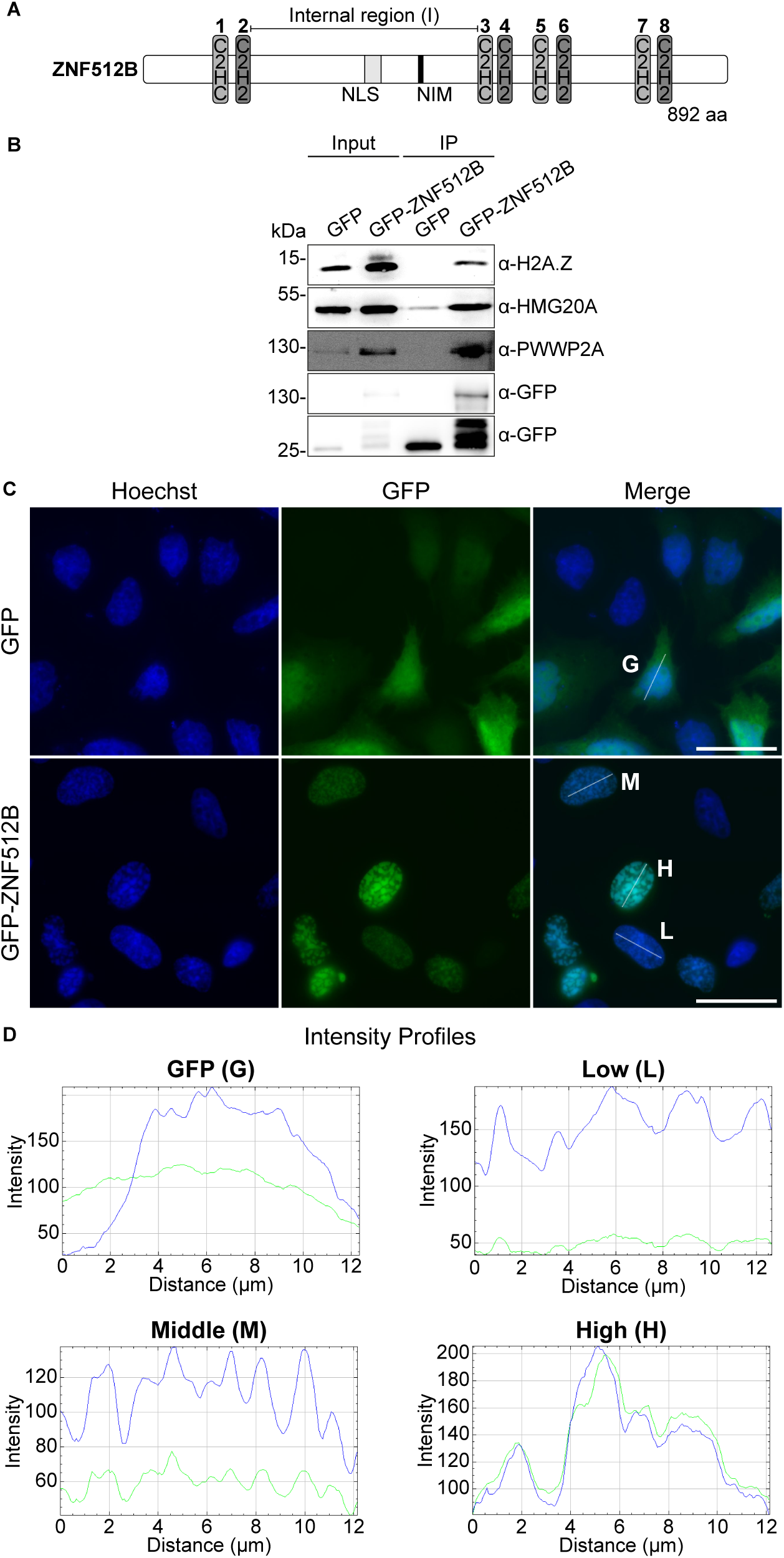
ZNF512B interacts with H2A.Z, HMG20A and PWWP2A and its overexpression leads to nuclear foci formation containing compacted chromatin. **(A)** Schematic representation of human ZNF512B protein with four pairs of atypical C2HC (light grey) and typical C2H2 (dark grey) ZFs and an internal (I) region containing an NLS (grey) and a putative NuRD interaction motif (NIM) (black). **(B)** Immunoblots of nuclear extracts from HeLaK cells transiently expressing GFP or GFP-ZNF512B after pull down with GFP-TRAP beads detecting H2A.Z and its associated HMG20A and PWWP2A proteins. **(C)** Immunofluorescence microscopy of HeLaK cells expressing GFP or GFP-ZNF512B (green) co-stained with Hoechst (DNA, blue). Scale bar = 20 µm. **(D)** Intensity profiles of nuclear areas from IF pictures in **(C)** (see lines) depicting DNA (blue) within cells expressing GFP (G), low (L), medium (M) or high (H) levels of GFP-ZNF512B (green).

First, we transfected HeLaK cells with plasmids encoding either GFP or a GFP-ZNF512B fusion. We note in passing that we were not able to generate stably expressing GFP-ZNF512B HeLaK cells, indicating that GFP-ZNF512B overexpression is – after some time – detrimental to cells. GFP-ZNF512B efficiently pulled down H2A.Z, PWWP2A and HMG20A, further confirming an association between these proteins (Figure 1B).

Immunofluorescence (IF) microscopy revealed that GFP-ZNF512B forms nuclear foci, whose sizes directly correlated with the level of GFP-ZNF512B expression (Figure 1C, D). Interestingly, DNA co-staining revealed a striking co-localization of chromatin to these foci, suggesting that GFP-ZNF512B overexpression is correlated with some sort of induced DNA/chromatin compaction/condensation. This surprising and unusual phenotype was independent of the location of the GFP tag (Supplementary Figure S1B) and the cell type used for transfections (Supplementary Figure S1C).

Although histone posttranslational modification (PTM) patterns in the nuclei were overall similar to GFP expressing cells, a decrease in H3K4me3 and H2A.Zac as well as a reciprocal increase in H3K27me3 signals at the sites of aggregation was observed in a majority of cases (Supplementary Figure S1D). Further, GFP-ZNF512B induced DNA/chromatin compaction foci did not represent prophase condensation spots, as the foci were not marked with the mitotic modification H3 serine 10 phosphorylation (H3S10ph) (Supplementary Figure S1E). GFP-ZNF512B chromatin foci were distinct from senescence-associated heterochromatin foci (SAHFs) (56), as revealed by the absence of SA-β-galactosidase staining (Supplementary Figure S1F). We also note that long-term expression of GFP-ZNF512B resulted in the death of cells, likely by apoptosis, a phenotype observed previously (57). Due to the timing, we conclude that apoptosis is not actively initiated by ZNF512B but more likely a consequence of the persistent chromatin aggregation phenotype.

These data suggest that high levels of ZNF512B cause protein as well as DNA/chromatin aggregation within the nucleus.

### ZNF512B interacts with the NuRD complex via an internal NuRD interaction motif (NIM)

Closer visual inspection of the amino acid composition of the human ZNF512B protein revealed, besides its eight ZF domains, a sequence within its internal region (I) that resembles the known NuRD interaction motif (NIM), which is also called the FOG-1-like motif (30) (Figure 2A). This ‘RRKQxxP’ motif (where ‘x’ is any amino acid) was first identified in the N-terminus of FOG-1 (30), and later in several other chromatin-regulating proteins, such as SALL1 (58), SALL4 (59), BCL11A (60) and ZNF827 (61), and mediates a direct interaction with the RBBP4 (RbAp48) subunit of the NuRD complex (38). We therefore wondered whether ZNF512B is also able to interact with NuRD via this motif and whether such a ZNF512B-NuRD repressor complex might be recruited to ZNF512B bound chromatin, leading to its deacetylation, remodelling and compaction. To experimentally test these two ideas, we transiently expressed GFP alone as control or GFP-ZNF512B in HeLaK cells and performed IPs with GFP-TRAP beads of DNaseI digested cell extracts. Immunoblots of the eluted material with antibodies against several NuRD components confirmed that ZNF512B is indeed a novel NuRD interacting protein (Figure 2B).

**Figure 2:**
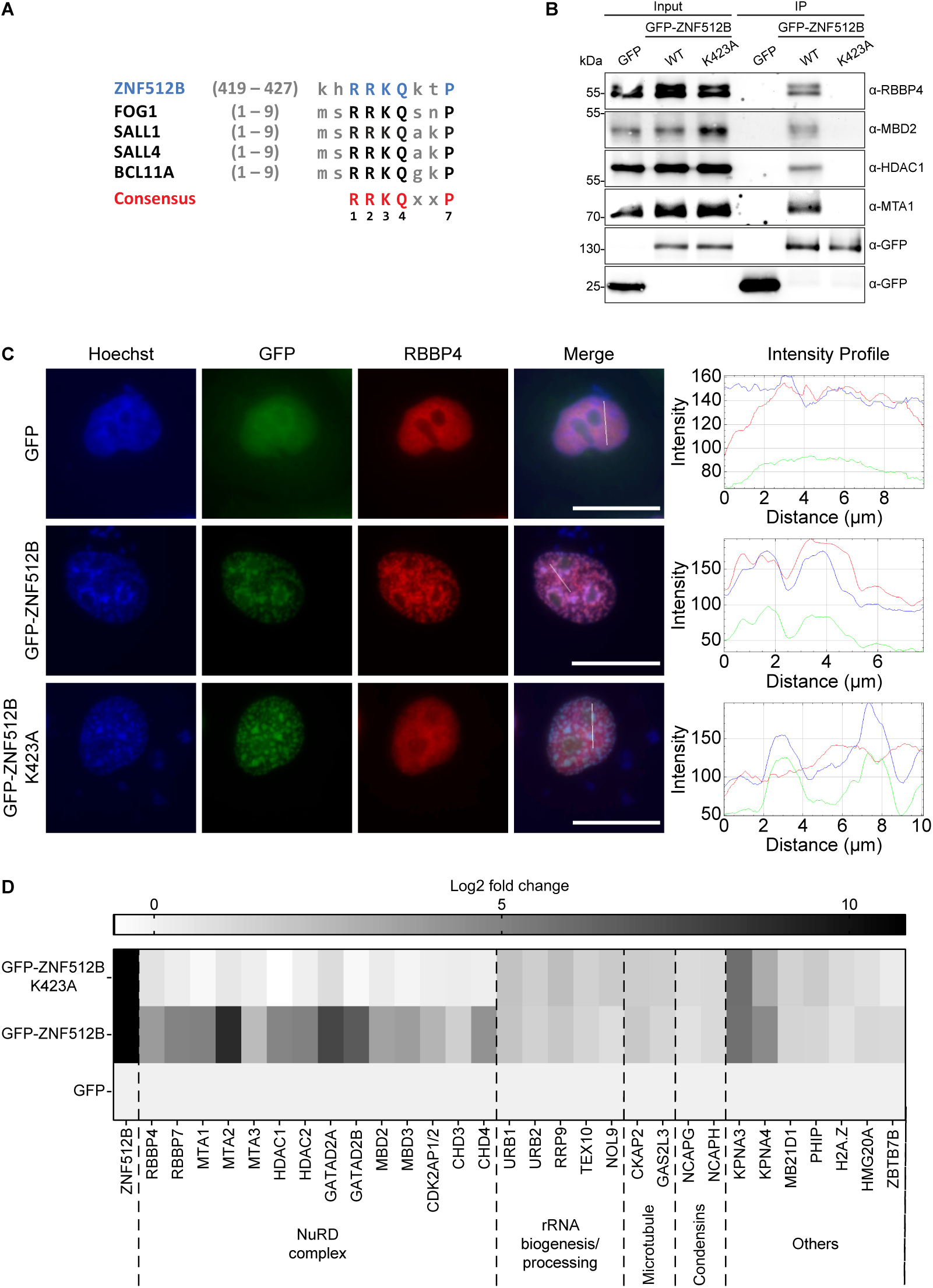
ZNF512B interacts with the complete NuRD complex via its NIM. **(A)** Alignment of ZNF512B’s putative NIM with N-terminal NIM sequences of other known NuRD binding proteins. **(B**) Immunoblots of cell extracts from HeLaK cells transiently expressing GFP, GFP-ZNF512B or GFP-ZNF512B_K423A mutant after pull down with GFP-TRAP beads detecting NuRD complex proteins. **(C**) Left: Immunofluorescence microscopy of HeLaK cells expressing GFP, GFP-ZNF512B or GFP-ZNF512B_K423A (green) co-stained with Hoechst (DNA, blue) and antibody against RBBP4 (red). Scale bar = 20 µm. Right: Intensity profile of nuclear areas (see lines, left) depicting DNA (blue), GFP constructs (green) and RBBP4 (red) fluorescence (see Supplementary Figure S2A, B for IF microscopy co-stains with NuRD members MTA1 and CHD4). **(D)** Heatmap of label-free mass spectrometry quantification of interactors with either GFP-ZNF512B or GFP-ZNF512B_K423A compared to GFP. Shown is the log2-fold average enrichment of two independent experiments with four technical replicates per sample each (see Supplementary Figure S2C-E for representative volcano plots).

Next, we verified this interaction by IF microscopy. Staining of GFP or GFP-ZNF512B expressing HeLaK cells with antibodies against the NuRD subunits RBBP4, MTA1 and CHD4 revealed a clear co-localization of these proteins at nuclear GFP-ZNF512B foci (Figure 2C, Supplementary Figure S2A, B). Having demonstrated the ZNF512B interaction with NuRD complex members, we next investigated whether the putative NIM in ZNF512B is responsible for mediating this interaction. We mutated the highly conserved lysine 423 in GFP-ZNF512B (Figure 2A) to alanine (K423A), because the corresponding residue in FOG-1 was shown to be required for NuRD binding (30). The GFP-ZNF512B_K423A mutant was transiently expressed in the nuclei of HeLaK cells but was unable to pull down NuRD proteins, in contrast to wild type GFP-ZNF512B (Figure 2B). Accordingly, the NuRD members RBBP4, MTA1 and CHD4 did not co-localize with GFP-ZNF512B_K423A (Figure 2C, Supplementary Figure S2A, B), demonstrating that the NIM of ZNF512B is necessary for NuRD interaction. Surprisingly, chromatin foci were still formed when overexpressing GFP-ZNF512B_K423A, indicating that this form of DNA/chromatin compaction is triggered independently of NuRD binding.

We further verified this NIM-dependent NuRD interaction by performing lf-qMS after GFP-TRAP IP of GFP, GFP-ZNF512B or GFP-ZNF512B_K423A. NIM-dependent binding of ZNF512B to all known NuRD complex members was observed (Figure 2D, Supplementary Figure S2C-E). Interestingly, we also detected NIM-independent binding of ZNF512B to several other proteins that are involved in nuclear processes, such as ribosome biogenesis. We biochemically confirmed the interaction of overexpressed GFP-ZNF512B and GFP-ZNF512B_K423A with Karyopherin Subunit Alpha 4 (KPNA4), a chaperone that also binds protein aggregates (62) (Supplementary Figure 2F), showing the validity of our approach.

In summary, we identified a FOG-1 resembling internal NIM sequence in ZNF512B that mediates an interaction with the NuRD complex but is not required for ZNF512B-induced chromatin compaction.

### The ZNF512B NIM directly interacts with RBBP4

Previously, FOG-1 has been demonstrated to bind to the RBBP4 subunit of the NuRD complex via its N-terminal NIM (38). We therefore wondered whether ZNF512B is also a direct interaction partner of RBBP4. Hence, we determined the X-ray crystal structure of an RBBP4-ZNF512B complex. As previously (63), we expressed His-tagged full-length RBBP4 in Sf9 insect cells, purified the protein and reconstituted the RBBP4-ZNF512B complex by combining RBBP4 with a synthetic peptide comprising the ZNF512B NIM (residues 419-430). We determined the X-ray crystal structure of this complex to an overall resolution of 2.2 Å (Supplementary Information). Two copies of the RBBP4-ZNF512B complex were found in the asymmetric unit; these had essentially identical conformations and we focus on one of these.

In the structure, RBBP4 forms the barrel-shaped β-propellor fold with a prominent acidic cavity at one of the apical surfaces (Figure 3A). We were able to model residues 10–410 into the electron density map, with the exception of a loop encompassing residues 90–113. This loop is absent in other structures of RBBP4 (*e.g.* (38)) and is likely to be highly dynamic; indeed, the AlphaFold2 model for human RBBP4 displays low confidence for the same region. The backbone conformation of RBBP4 in this complex also closely resembles that of the same protein in the RBBP4-FOG-1 complex (PDB: 2xu7), with a backbone RMSD of 0.29 Å (Supplementary Figure 3A).

**Figure 3:**
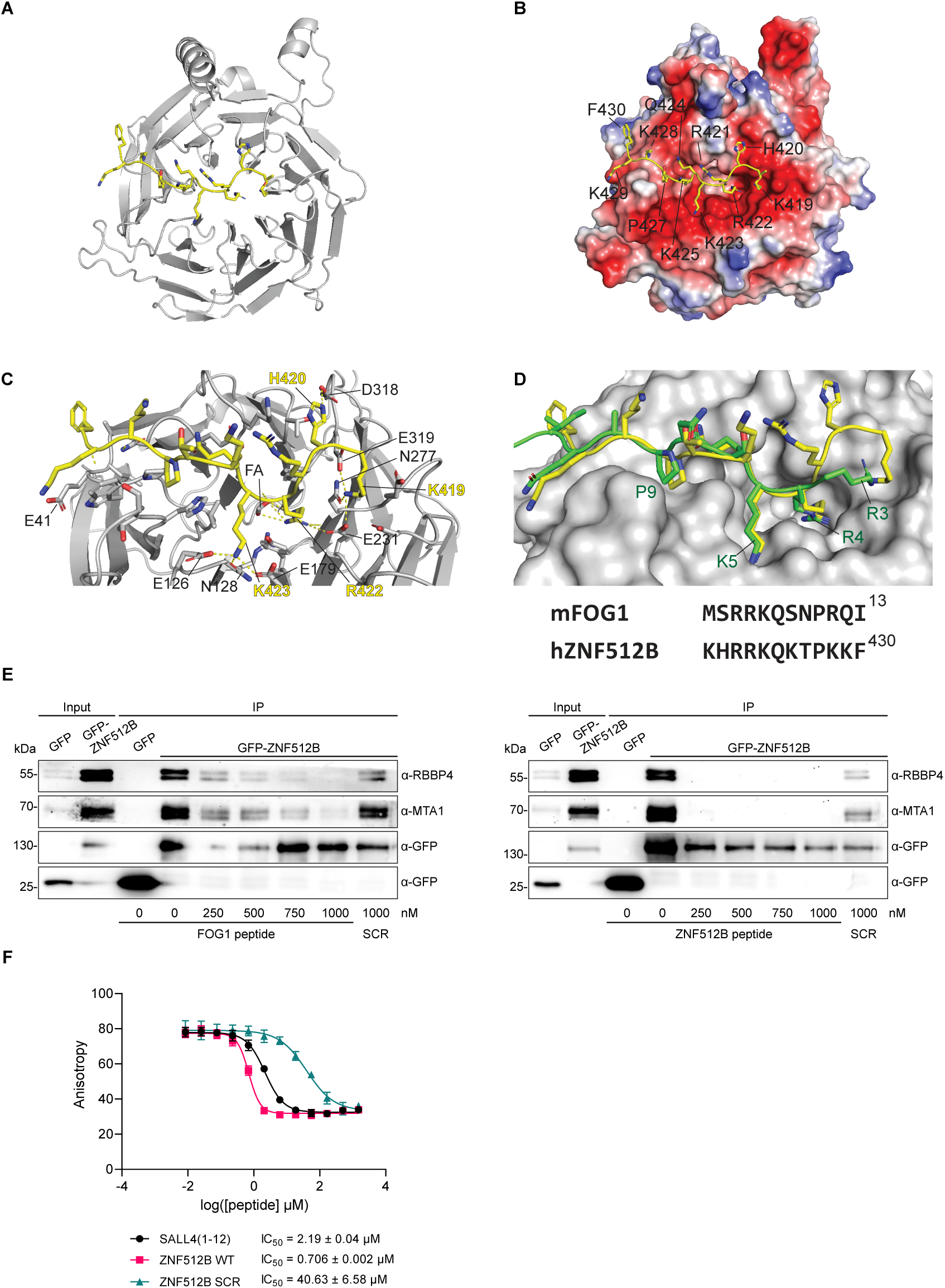
ZNF512B directly binds RBBP4 with high affinity. **(A)** Ribbon diagram of the RBBP4-ZBF512B complex. RBBP4 is shown in grey and ZNF512B in yellow. **(B)** Same as (A) but with RBBP4 shown as an electrostatic potential surface (created in Pymol). **(C)** Close-up of polar interactions (dashed lines) made between RBBP4 and ZNF512B. Residue numbers are indicated in black for RBBP4 and yellow for ZNF512B. **(D)** Overlay of the structure of FOG-1(1–12) (PDB: 2XU7, green) bound to RBBP4 (grey) (38) with ZNF512B (yellow). Key residues in FOG-1 are labelled. A sequence alignment is shown below. **(E)** Immunoblots of cell extracts from HeLaK cells transiently expressing GFP or GFP-ZNF512B mixed with increasing amounts of FOG-1 (left) or ZNF512B (right) NIM peptides or respective scrambled (SCR) controls before pull down with GFP-TRAP beads detecting NuRD complex members RBBP4 and MTA1. **(F)** Fluorescence polarization-based binding experiment of RBBP4. Competitive experiments of unlabeled SALL4(1-12), ZNF512B wild-type NIM and scrambled peptides displacing Ac-SALL4(2-12)-FAM (1 nM) from RBBP4 (1.25 µM).

The entire 11 residues of ZNF512B (419–430) can be modelled confidently in the structure, showing that the peptide binds in an extended conformation to the acidic surface of RBBP4 (Figure 3B). A total of 1660 Å^2^ of surface area is buried in the formation of the complex and a large number of electrostatic interactions are formed (Figure 3C). In general, residues in the C-terminal half of the ZNF512B peptide (Q424–F430) take up very similar conformations to the corresponding residues of FOG-1 in the RBBP4-FOG-1 structure (PDB: 2XU7, (38); Figure 3D). The sidechains of K425, T426 and K428 project into solvent, whereas Q424 and P427 pack into a broad uncharged groove. K429 lies on an acidic surface that is made from E41 and D74 in one of the two copies of the complex and E41 and E75 in the other; the RBBP4 backbone undergoes a localized reorientation to position either D74 or E75 near the K429 sidechain.

Similarly, the sidechain positions of ZNF512B residues R422 and K423 lie in the same negatively charged pockets as the corresponding residues in FOG-1 and form similar interactions seen in FOG1 (Figure 3B-D). Thus, K423 interacts with the sidechains of E126, N128 and E179, whereas R422 forms an ion pair with E231 and the backbone carbonyl of N277. We observed additional electron density in the R422 pocket and could fit this to a molecule of formic acid, which was present in the crystallization buffer and makes electrostatic interactions with the R422 guanidino group.

Unexpectedly, significant differences were observed in the N-terminal segment of the peptide. In the RBBP4:FOG-1 structure (Figure 3D), the sidechain of arginine at position 3 lies on an acidic ‘shelf’, making electrostatic interactions with the sidechains of E231, N277 and E319. However, a 130° rotation of the backbone psi angle of R421 in ZNF512B reorients the sidechain of this residue so that it faces away from RBBP4 and towards the solvent. Instead, the sidechain amino group of K419 occupies the corresponding shelf (Figure 3B) and forms an essentially identical set of interactions with RBBP4 – with the sidechains of E231, N277 and E319 (Figure 3C). Furthermore, the sidechain of H420 packs against the acidic pocket, interacting with D318 and contributing to this reorientation of the peptide relative to the conformation observed for FOG-1.

Given this difference in binding mode, we used pull-downs to ask whether ZNF512B and FOG-1 might have different binding affinities to RBBP4. We used GFP-TRAP beads to capture GFP-ZNF512B from HeLaK cell extracts in the absence or presence of increasing amounts of either FOG-1 or ZNF512B NIM peptides, detecting the precipitated NuRD components MTA1 and RBBP4 in immunoblots (Figure 3E). Less than 250 nM of the ZNF512B peptide prevented binding of GFP-ZNF512B to MTA1 and RBBP4, whereas these interactions were not completely competed with >750 nM FOG-1 peptide. This result suggests that the NIM in ZNF512B has a higher binding affinity for RBBP4 than the corresponding motif in FOG-1. Furthermore, competitive Fluorescence Polarization (FP)-based experiments using the SALL4 tracer, which shares the same sequence as FOG-1 but one amino, reveal that the ZNF512B’s NIM peptide can efficiently displace SALL4 tracer (IC_50_ = 0.706 ± 0.002 µM), while its scrambled version displayed a > 65-fold lower interaction (IC_50_ = 40.63 ± 6.58 µM) (Figure 3F).

In conclusion, the NIM in ZNF512B binds to RBBP4 in a manner that is partially distinct from the mode of binding used by FOG-1 in its interaction with this NuRD component.

### Distinct amino acids in the ZNF512B NIM are required and necessary for NuRD binding

The crystal structure of RBBP4 binding to the internal NIM of ZNF512B revealed a network of interactions between the two proteins. To independently test the requirement of each amino acid within this motif in a cellular context, we carried out an alanine mutagenesis scan, generating a set of GFP-ZNF512B mutants that each carried a point mutation of one residue in the NIM. As expected, R422, K423 and P427 of the FOG-1-predicted ‘RRKQxxP’ motif were essential for NuRD binding (Figure 4A). Similar results were obtained when looking at the co-localization of all GFP-ZNF512B alanine mutants and RBBP4 in IF microscopy (Figure 4B and Supplementary Figure S4A).

**Figure 4:**
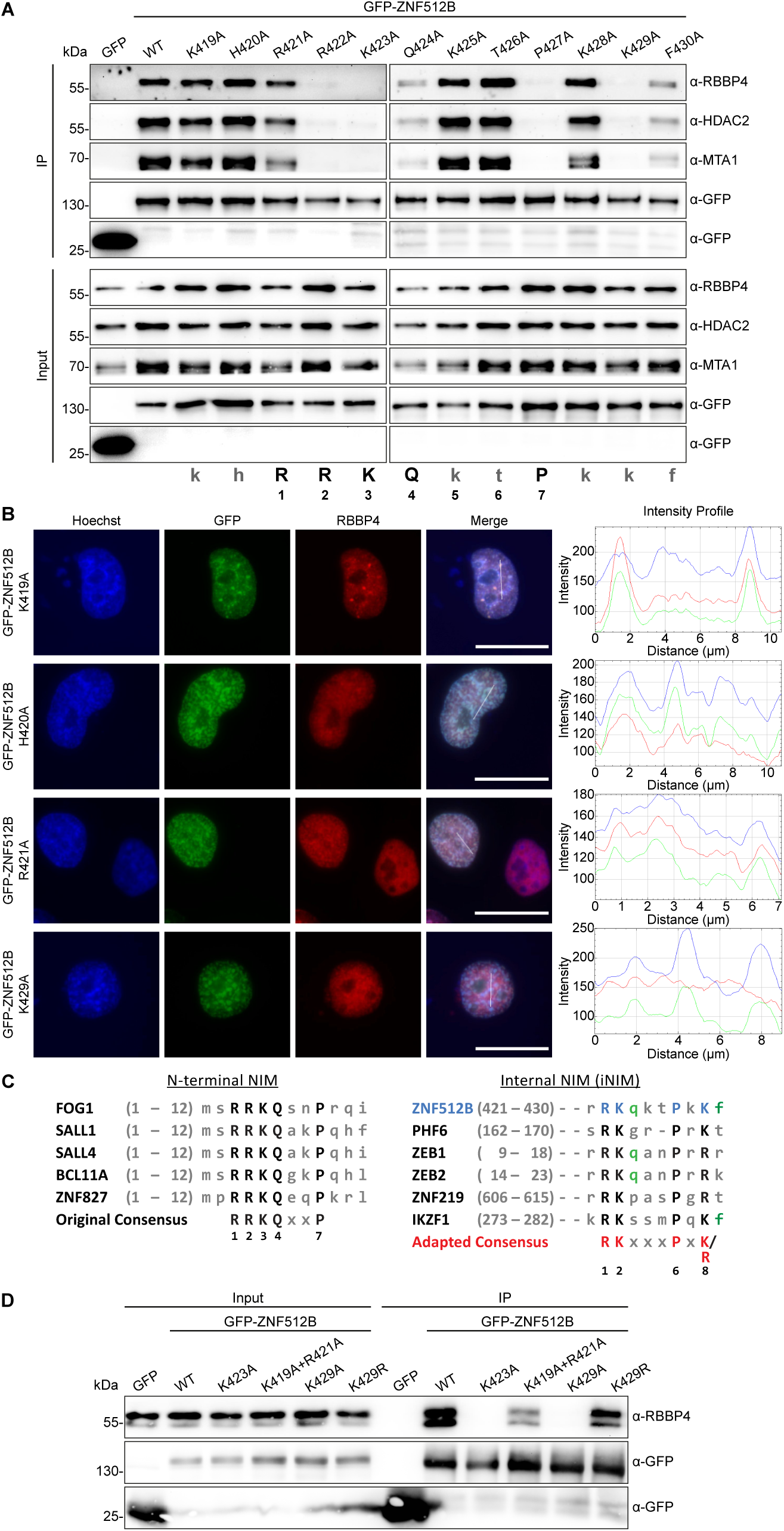
A mutational screen reveals a functional internal NIM (iNIM) sequence in ZNF512B. **(A)** Immunoblots of cell extracts from HeLaK cells transiently expressing GFP and GFP-ZNF512B NIM alanine exchange mutants after pull down with GFP-TRAP beads detecting NuRD complex members RBBP4, MTA1 and HDAC2. NIM consensus sequence is shown at the bottom. **(B)** Left: Immunofluorescence microscopy of HeLaK cells expressing GFP and some GFP-ZNF512B NIM alanine exchange mutants (green) co-stained with Hoechst (DNA, blue) and anti-RBBP4 antibody (red). Scale bar = 20 µm. Right: Intensity profile of nuclear areas (see lines, left) depicting DNA (blue), GFP constructs (green) and RBBP4 (red) fluorescence. See Supplementary Figure S4A for IF of all mutants. **(C)** Alignments of original, N-terminal (left) and internal NIM (iNIM, right) consensus sequences. **(D)** Immunoblots of cell extracts from HeLaK cells transiently expressing GFP, GFP-ZNF512B or GFP-ZNF512B_K429R mutant after pull down with GFP-TRAP beads detecting NuRD complex member RBBP4. See Supplementary Figure S4B for IF of GFP-ZNF512B_K429R.

Similar to a previously published FOG-1 interaction screen (30), we found that the arginine (R421) in the 1^st^ position within the consensus motif was not required for NuRD binding and that the glutamine (Q424) in the 4^th^ position was only minimally required (Figure 4A). In agreement with our RBBP4-ZNF512B crystal structure, we found K429 to be essential for and F430 to partially contribute to NuRD interaction (Figure 4A). Similar results were obtained when looking at the co-localization of all GFP-ZNF512B alanine mutants and RBBP4 in IF microscopy (Figure 4B and Supplementary Figure S4A).

Taking these data into account, we wondered whether the so far published NIM consensus sequence needs to be adapted. We compared the up-to-now known N-terminally located NIMs from FOG-1, SALL1, SALL4, BCL11 and ZNF827 (59,64-66) with the internally located sequences in PHF6 (67) and ZNF512B. On the basis of this comparison, we propose the existence of two distinct RBBP4/NuRD interaction consensus motifs: i) an N-terminal ‘RRKQxxP’ and ii) an internally localized ‘RKxxxPxK’ sequence that we termed internal NIM (iNIM) (Figure 4C). Based on this prediction, we used the ScanProsite tool from Expasy to search for human proteins containing an iNIM and found several proteins that in the past have been found to be somehow connected to NuRD, including ZEB2 (68) and IKZF1 (69) (Figure 4C). Because these two examples contain an arginine instead of a lysine residue at the 8^th^ position within the motif, which was shown to be necessary for NuRD interaction, we wondered whether an arginine residue at this site is also sufficient for RBBP4 interaction. To test this idea, we exchanged the according K429 of ZNF512B for arginine and found that this mutant retains binding to NuRD, as shown by IP (Figure 4D) and IF microscopy (Supplementary Figure S4B) experiments. Taking PHF6’s iNIM sequence into account, these data support the existence of a novel iNIM with the consensus sequence ‘RK(x)xxPxK/R’.

In summary, we have identified a functional internal RBBP4/NuRD-interaction consensus motif in ZNF512B that is also present in several other proteins.

### ZNF512B overexpression and depletion affects gene expression in a NuRD-dependent and -independent manner

Having demonstrated ZNF512B binding to NuRD via an iNIM, we next wondered whether ZNF512B has any role in regulating gene transcription and whether such a process depends on NuRD binding. First, we transiently overexpressed GFP, GFP-ZNF512B (NuRD binding) or GFP-ZNF512B_K423A (loss of NuRD binding) in HeLaK cells, isolated RNA two days after transfection and performed high-throughput sequencing (RNA-seq) (Supplementary Figure S5A). We found only few genes to be differentially expressed, with around 500 or 350 genes to be significantly deregulated upon GFP-ZNF512B or GFP-ZNF512B_K423A overexpression in comparison to GFP expressing control cells, respectively (Figure 5A). Of those genes, 229 genes were exclusively deregulated in GFP-ZNF512B overexpressing cells, while 68 genes were exclusively deregulated upon GFP-ZNF512B_K423A overexpression (Supplementary Figure S5B). Most of the commonly and GFP-ZNF512B_K423A exclusively deregulated genes were downregulated, while, surprisingly, the majority of GFP-ZNF512B exclusively deregulated genes was upregulated (Figure 5B). Such upregulated genes were mainly expressed at extremely low levels or not expressed at all in GFP control cells (Supplementary Figure S5C). Over representation analysis (ORA) and gene set enrichment analysis (GSEA) for both Gene Ontology (GO) and Kyoto Encyclopedia of Genes and Genomes (KEGG) databases indicate that shared deregulated genes belong to several different biological processes and molecular functions, not allowing us to draw any clear biological link between ZNF512B and a precise process (Supplementary Figure S5D). These data suggest that ZNF512B is involved in transcriptional regulation, in a NuRD-dependent and -independent manner. As we used an artificial over-expression system so far, we next wondered whether the endogenous protein also plays a role in controlling gene expression. We therefore depleted endogenous ZNF512B in HeLaK cells by siRNA-mediated knock-down (KD). We observed an efficient reduction of ZNF512B mRNA and protein two days after transfection using RT-qPCR (Supplementary Figure S5E), immunoblots (Figure 5C) and IF microscopy (Supplementary Figure S5F). We then performed RNA-seq (two biological replicates) on these samples (Supplementary Figure S5G). We found 18 genes to be up- and 8 to be down-regulated significantly upon ZNF512B depletion (Figure 5D), suggesting that ZNF512B is indeed involved in the transcriptional regulation of a subset of genes. ORA and GSEA analyses for GO and KEGG databases revealed that deregulated genes upon depletion of ZNF512B are mainly involved in RNA related processes (Supplementary Figure S5H). To determine direct target genes of ZNF512B that are repressed in a NuRD-dependent manner, we compared the RNA-seq data sets of all downregulated genes upon GFP-ZNF512B overexpression with upregulated genes in ZNF512B KD cells (Figure 5E). We detected only nine genes to be commonly deregulated in such a manner (Figure 5F, Supplementary Figure 5I). In summary, ZNF512B is involved in the regulation of gene expression.

**Figure 5:**
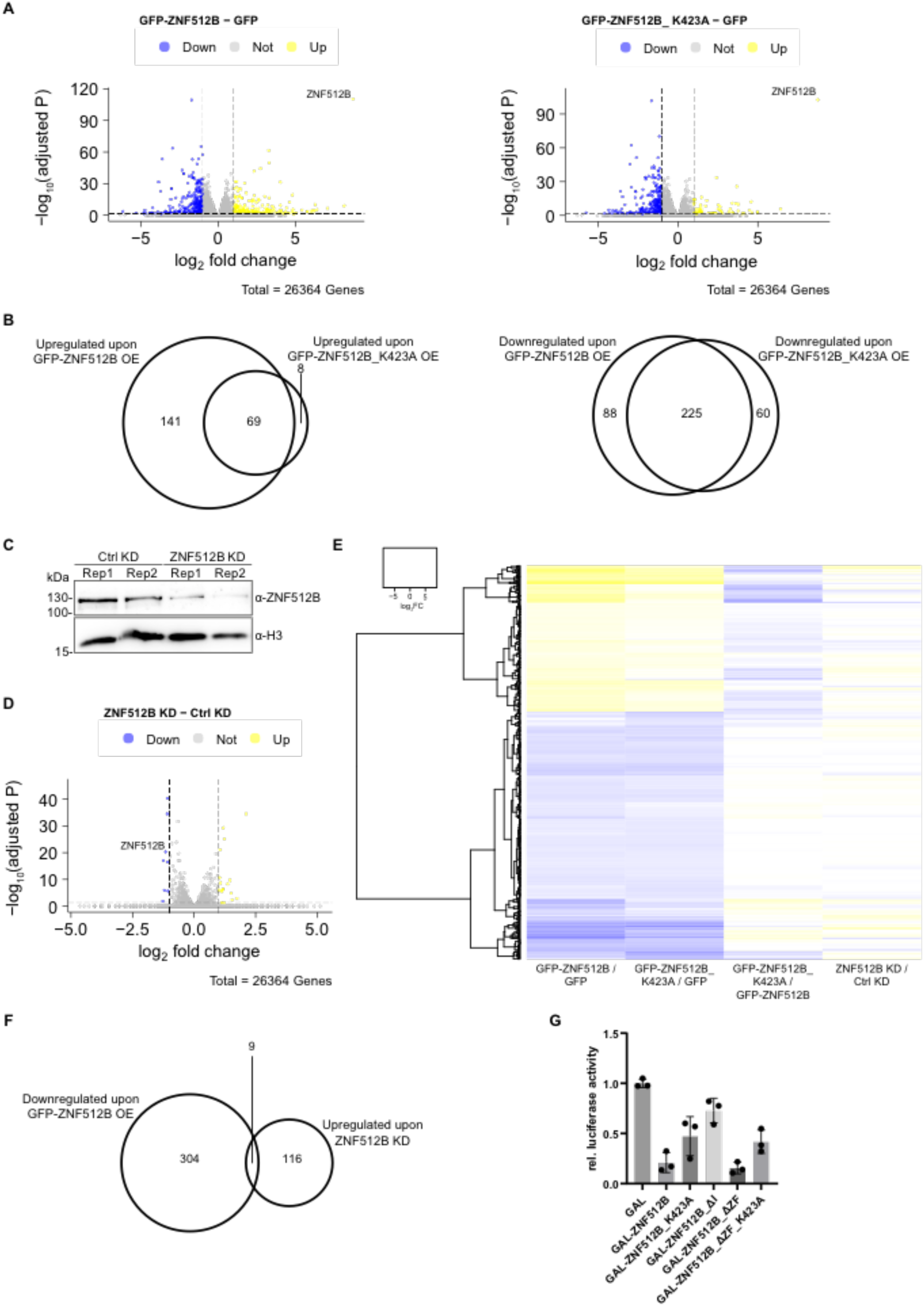
Overexpression and depletion of ZNF512B affects gene expression in NuRD-dependent and -independent manner. **(A)** Volcano plots showing significantly deregulated genes in GFP-ZNF512B or GFP-ZNF512B_K423A expressing cells compared to GFP control. Depicted is log2 fold change and adjusted p-values. **(B)** Euler diagram depicting the overlap of significantly up- or downregulated genes (log2 FC > 1 or < -1 and adjusted p-value < 0.05) upon GFP-ZNF512B or GFP-ZNF512B_K523A overexpression compared to GFP control. **(C)** Immunoblot of nuclear extracts from two HeLaK cell replicates (Rep) two days after transfection with control (Ctrl) or ZNF512B-specific siRNA pools stained with ZNF512B or H3 (loading control) antibodies to verify ZNF512B knock-down (KD) efficiency. **(D)** Volcano plot showing mean of differentially deregulated genes in ZNF512B KD cells. Depicted is log2-fold change of the mean of two biological RNA-seq replicates compared to control KD. **(E)** Heatmap of deregulated genes upon GFP-ZNF512B or GFP-ZNF512B_K423A overexpression or ZNF512B depletion. Shown is the log2-fold change of GFP-ZNF512B / GFP, GFP-ZNF512B_K423A / GFP, GFP-ZNF512B / GFP-ZNF512B_K423A or ZNF512B / Ctrl KD comparison. **(F)** Euler diagram showing the overlap of upregulated genes (log2 FC > 0.5 and adjusted p-value < 0.05) upon ZNF512B KD and downregulated genes (log2 FC < -1 and adjusted p-value < 0.05) upon GFP-ZNF512B overexpression. **(G)** Reporter assay of HEK293T cells transiently transfected with luciferase-reporter and GAL control or different GAL-ZNF512B wild type or deletion/mutation plasmids. SD = 3.

Lastly, we utilized a luciferase-based reporter assay in HEK293T cells to determine whether promoter recruitment of ZNF512B and its deletions can affect transcription in a NuRD-dependent or -independent manner. Interestingly, ZNF512B strongly repressed reporter gene expression (Figure 5G), suggesting that it might rather be involved in the inhibition than the activation of gene transcription. To clarify whether this effect is due to its interaction with the NuRD complex, we utilized several ZNF512B deletion or mutation constructs. Surprisingly, loss of NuRD binding following K423A mutation still resulted in 50% repression, suggesting that NuRD interaction is only partially responsible for the inhibitory effect. On the other hand, deletion of the entire internal region completely abolished the repression effect, hinting at the existence of an additional repressive region within ZNF512B. Likewise, expression of the internal region alone without ZFs resulted in strong repression similar to the full-length ZNF512B protein, whereas the additional K423A mutation again reduced the repression ability to 50%.

In conclusion, ZNF512B acts as a transcriptional repressor mediated by its internal region and regulates a subset of genes in both NuRD-dependent and -independent ways.

### Size and formation of nuclear ZNF512B/chromatin foci depend on the number of ZFs

Finally, we set out to determine how high levels of ZNF512B protein affect chromatin structure and gene expression pattern without any NuRD complex involvement. We generated several GFP-ZNF512B deletion constructs, which were transiently expressed in HeLaK cells, and the degree of DNA compaction was visualized by Hoechst staining. As expected, GFP-ZNF512B overexpression led to the formation of nuclear protein aggregates and large DNA foci (Figure 6A). When all ZFs were deleted and only the internal region containing the iNIM was expressed, both protein and DNA aggregation was absent (Figure 6B), suggesting that the ZF domains are responsible for this phenotype. Indeed, expression of all ZFs without the internal region resulted in protein aggregation and DNA compaction/condensation similar to wild type GFP-ZNF512B (Figure 6C). Surprisingly, stepwise deletion of ZF pairs correspondingly reduced the sizes of protein aggregates and DNA foci (Figure 6D, E). As the severity of the ZNF512B and chromatin aggregation phenotype depends on the number of ZFs, we speculate that they participate in i) oligomerization, explaining the protein aggregation phenotype and the interaction with KPNA3/4 proteins and ii) DNA binding, which mediates the observed chromatin compaction/condensation. To first test ZNF512B’s ability to di-/oligomerize we performed co-transfections with GFP- and FLAG-tagged ZNF512B constructs, followed by GFP-TRAP IPs and FLAG immunoblotting. These experiments showed that ZNF512B is able to form oligomers (Figure 6F).

**Figure 6:**
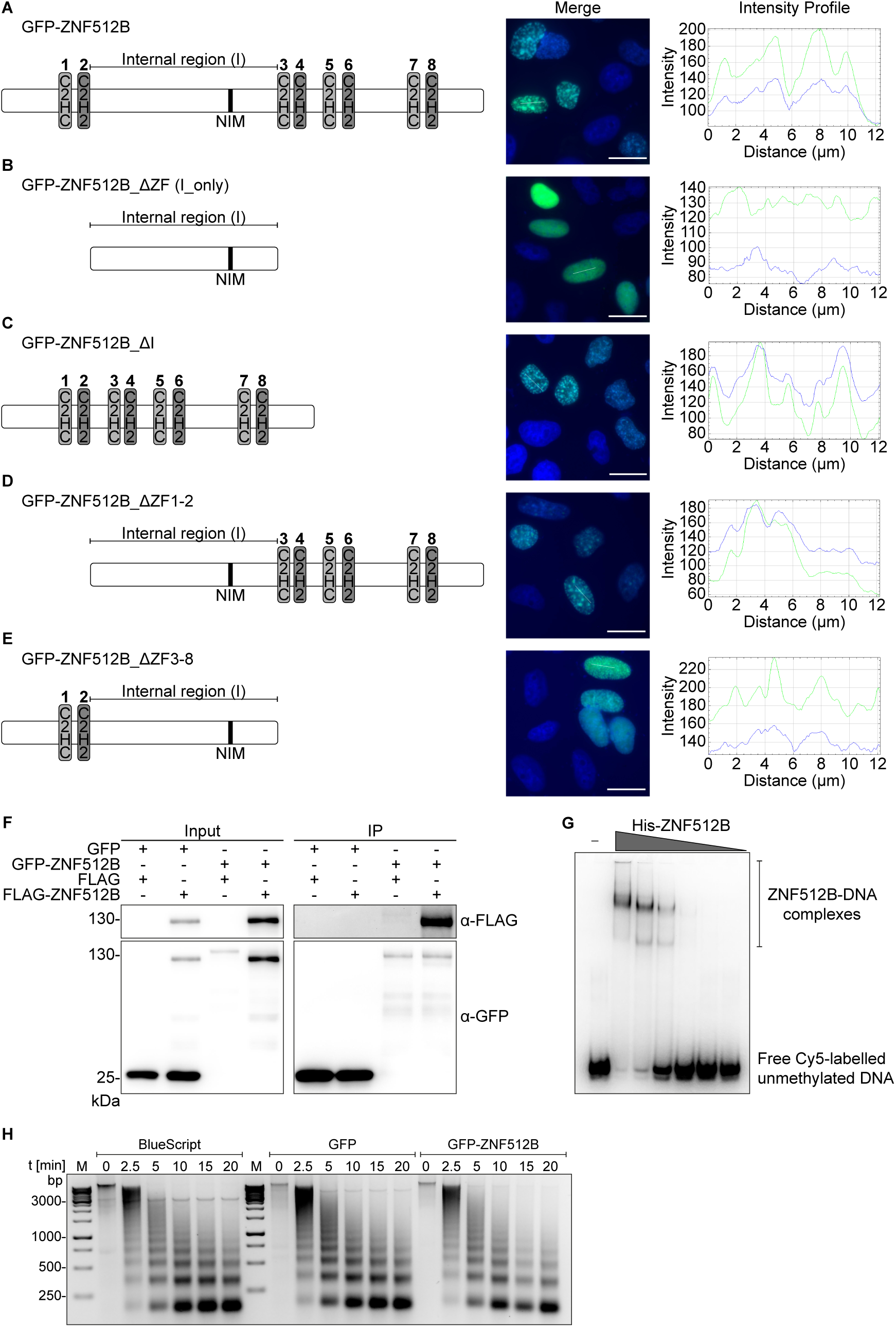
The number of ZNF512B ZFs determines chromatin compaction severity. Left: Schematic depiction of used constructs. Middle: IF microscopy of HeLaK cells expressing **(A)** GFP-ZNF512B, **(B)** GFP-ZNF512B_ΔZF (internal domain with NIM only), **(C)** GFP-ZNF512B_ΔI (only ZFs, without internal region), **(D)** GFP-ZNF512B_ΔZF1-2 or **(E)** GFP-ZNF512B_ΔZF3-8 (green) co-stained with Hoechst (DNA, blue). Scale bar = 20 µm. Right: Intensity profile of nuclear areas (see lines, left) depicting DNA (blue), GFP constructs (green) and RBBP4 (red) fluorescence. **(F)** Immunoblots of cell extracts from HeLaK cells expressing GFP or GFP-ZNF512B together with FLAG or FLAG-ZNF512B after pull down with GFP-TRAP beads detecting GFP (IP-control) and FLAF-ZNF512B. **(G)** Electrophoretic Mobility Shift Assay (EMSA) of Cy5-labelled DNA together with increasing amounts of purified recombinant His-ZNF512B protein**. (H)** Agarose gel of MNase (different-time points) digested chromatin isolated from HeLaK cells (control; Bluescript vector transfection; left) expressing GFP (middle) or GFP-ZNF512B (right).

Electrophoretic Mobility Shift Assays (EMSAs) with purified recombinant His-ZNF512B protein (Supplementary Figure S6A) also show that ZNF512B is a DNA-binding protein (Figure 6G), independent of DNA methylation status (Supplementary Figure S6B). We hypothesize that ZNF512B might oligomerize and bring together chromatin domains by simultaneously binding to DNA. In such a case, ZNF512B-triggered chromatin aggregation would not result in a direct change in nucleosome occupancy, but rather brings chromatin domains into closer contact. This idea is supported by a time-course MNase digestion experiment (Figure S6H), which showed no dramatic changes in chromatin accessibility after GFP-ZNF512B overexpression.

These data indicate that ZNF512B binds DNA and oligomerizes, and that its ZFs are required for chromatin aggregation independent of the internal region or NuRD recruitment.

## Discussion

We have identified ZNF512B as novel RBBP4-mediated NuRD complex interacting protein, whose overexpression leads to chromatin aggregation independent of NuRD binding.

Previously, we have found ZNF512B to be associated with H2A.Z and its binding proteins PWWP2A and HMG20A in diverse mass spectrometry screens (16,19,20,34) and could confirm these associations in IP and lf-qMS experiments. Now, we report that ZNF512B is also a direct interactor of the NuRD complex via the RBBP4 subunit. There have been previous hints about ZNF512B possibly being a binding partner of NuRD from several mass spectrometry-based surveys. These studies identified ZNF512B to be associated with the NuRD complex members MBD3 and GATAD2B (26,70), MTA1/2/3 (71) and HDAC1/2 but not with other HDAC family proteins (72). However, how exactly and via with which member(s) ZNF512B interacts with the NuRD complex was not investigated. We now demonstrate that the ZNF512B-NuRD interaction is mediated by ZNF512B’s internal NIM (iNIM). This facilitates a direct interaction that has a higher affinity than the RBBP4-FOG-1 or RBBP4-SALL1 interactions.

This internal NIM consensus sequence differs from the published ‘RRKQxxP’ motif that has previously been identified in the N-termini of e.g. FOG-1 (30), SALL1 (58), SALL4 (59), BCL11A (60), ZNF296 (73) and ZNF827 (61). Similar to the published RBBP4-ZNF827 interaction (61), we also found the first arginine of the motif to be dispensable for this interaction. Further, the conserved glutamine (Q) residue at the 4^th^ position of the motif does not appear to be required for NuRD binding. Notably, ZNF296, another NuRD interaction protein that also pulls down ZNF512B (73), even contains a slightly altered N-terminal consensus sequence ‘RRKAxxxP’ with an alanine instead of a glutamine at the 4^th^ position, suggesting that this particular residue might not be as important for NuRD complex binding as yet thought. Interestingly, the last basic residue (8^th^ position) of the iNIM ‘RK(x)xxPxK/R’ sequence is required for NuRD binding. Using this sequence in a database search, we identified several proteins. Some of those have already been described to be associated with RBBP4 or the complete NuRD complex, such as PHF6 (67), ZNF219 (74), ZEB2 (68), CENP-A (75), IKZF1 (Ikaros) (69), IKZF2 (Helios) (76), L3MBTL2 (77), TBX2 (78) and TRPS1 (79), strengthening the notion that this is indeed a novel functional iNIM sequence. Yet, many other proteins, such as the forkhead transcription factor FOXJ1 (80), the zinc finger protein ZNF148 (ZBP-89) (81) and the transcription factor ZBTB20 (82), have so far – and to our knowledge – not been connected to NuRD biology. Future experiments will provide more insights into whether these proteins are indeed novel RBBP4 and NuRD interacting proteins. One can envision two scenarios: i) They bind NuRD exclusively in a cell type-, stimulation- or differentiation-specific manner, explaining why they have not been identified so far in published mass spectrometry screens, and ii) they are not able to bind NuRD due to other protein features surrounding the iNIM sequence, such as e.g. secondary and tertiary structures that bury the iNIM, posttranslational inhibitory modifications or adjacent ‘blocking’ amino acids or due to a competition with other interactors, that in turn prevent direct binding to RBBP4. It is worth mentioning that NIM sequences can be further adapted. The PR-SET domain-containing proteins PRDM3 and PRDM16 have been found to interact with the NuRD complex by directly binding to RBBP4 using an N-terminal ‘RsKxrarrK’ sequence, with lysine at the 4^th^ position being essential for this interaction and lysine at the 9^th^ position forming hydrophobic and polar contacts with Q41 of RBBP4 (83). Also, the Epstein-Barr virus protein BMRF1 binds RBBP4 via an internal ‘RqKqkhPkK’ sequence (84). These data suggest that a NIM sequence can be more flexible, with an amino acid inclusion between the arginine at the 1^st^ and lysine at the 2^nd^ position and possibly the replacement of proline at the 6^th^ position.

Although we demonstrate a clear interaction between ZNF512B and RBBP4, we are, interestingly, not able to pull down other complexes that contain RBBP4, such as the PRC2 complex. This seems to be a common trait for proteins that bind RBBP4 via a NIM since the targeted binding pocket is blocked when the protein is engaged with PRC2, as structural studies show (85,86).

One unexpected observation of our study was the formation of protein and DNA/chromatin aggregates upon overexpression of ZNF512B. Interestingly, we observed an increase of H3K27me3 and a decrease of H3K4me3 and H2A.Zac within these aggregates, suggesting that predominantly facultative heterochromatin with an exclusion of actively transcribed euchromatic regions was targeted by GFP-ZNF512B, when present in high concentrations. While we are aware of the non-physiological challenges of such a rather artificial overexpression system, we are convinced that it can also offer new insights into ZNF512B biology. Endogenous ZNF512B mRNA is expressed ubiquitously at low level, however, brain as well as some cancer tissues, show elevated ZNF512B gene expression. It will therefore be of interest to determine in future experiments, whether cells in these tissues also show changes in chromatin structures, when levels of ZNF512B protein are altered. So far, all our data point towards ZNF512B being able to di-/oligomerize and to bind DNA. In turn, these interactions result when a high amount of ZNF512B proteins is present, in large and dense nuclear aggregates. We discovered that the severity of this phenotype depends on the number of ZFs present in ZNF512B and not on its interaction with NuRD. Further, we hypothesize that this form of aggregation also fosters the interaction of ZNF512B with KPNA3/4 karyopherins that, besides being involved in nuclear import (87), also have been reported to serve as chaperones for protein aggregates (62). Moreover, karyopherins are often found to be deregulated or mutated in protein-aggregation-dependent neurodegenerative diseases, such as Amyotrophic Lateral Sclerosis (ALS) (88), a disease whose survival span has been, albeit still controversial, associated with one particular SNP (rs2275294 polymorphism) in the regulatory region of the human *znf512b* gene in several meta-studies (27-29,89-91). It will therefore be of interest to determine whether ZNF512B expression levels are altered in such a cohort of ALS patients, maybe even elevated leading to disease-promoting protein and DNA aggregation. On the other hand, we found ZNF512B to also bind several rRNA processing proteins, such as URB1/2 (92), RRP9 (93), TEX10 (94) and NOL9 (95). These findings implicate ZNF512B as being putatively involved in ribosomal RNA processing (a GO term we also found upon ZNF512B depletion), a process also often affected in patients with ALS (96,97). It is therefore similarly possible that cells of such patients containing a SNP in the regulatory region of the *znf512B* gene have reduced ZNF512B protein levels and altered ribosome biogenesis, a hypothesis that will be investigated in planned future studies.

Finally, how does ZNF512B contribute to H2A.Z biology? As H2A.Z has been involved in – among other DNA-based processes – transcriptional repression and activation (98), it is possible that H2A.Z mediates these conflicting activities, although not exclusively, by the recruitment of distinct chromatin-modifying proteins (18). Therefore, ZNF512B binding to H2A.Z-containing chromatin regions might recruit NuRD and its deacetylase and remodelling activities to regulatory genomic regions thereby affecting gene expression, as we have seen in reporter assays and RNA-seq of both ZNF512B overexpressing and depleted cells. However, as more genes are also deregulated in a NuRD-independent manner, we speculate that deregulation of transcription in the overexpression system might be a consequence of a combination of chromatin aggregation and, most importantly, the influence of a yet uncharacterized part of the internal region that has some sort of repressor activity. Interestingly, most differentially deregulated genes were upregulated in the GFP-ZNF512B overexpressing cells. We hypothesize that GFP-ZNF512B competes with other proteins for NuRD binding and, as it is present in extremely high concentrations (> 5,000 times higher than endogenous protein, as determined by RNA-seq), it might bind to a large amount of the nuclear, repressive NuRD complexes, delocalizing them from their distinct target genes, in turn leading to an upregulation of their expression. As other ZF proteins have been shown to be recruited to only one single target gene, such as ZFP568 (99), ZFP64 (100), ZNF410 (101) and ZNF558 (102), it is plausible to assume that ZNF512B might also have only a few or even a single target sites that we are not able to identify with the approach here. Future ChIP studies will shed light on this fascinating possibility.

In general, we have identified ZNF512B as an RBBP4-mediated NuRD binding protein that is involved in gene repression and that causes chromatin aggregation when present at elevated levels.

## Data Availability

Raw and processed RNA-seq files are deposited in GSE236639. The mass spectrometry proteomics data have been deposited to the ProteomeXchange Consortium via the PRIDE (103) partner repository with the dataset identifier PXD043366.

## Supplementary Data

Supplementary Data are available online.

## Acknowledgements

We thank all current and past members of the Hake team for practical help, support and ideas, especially Sonja Sahner, Carlotta Kreienbaum, Charlotte Floh and Jonas Schauss.

## Author contributions

TMW and SBH conceived of this study. TMW cloned all constructs, performed IPs, IFs, peptide competition and RNA-seq with the help of LP, SES, ND and FD. JoLe and HN generated recombinant FLAG-ZNF512B protein in insect cells and performed cEMSAs and luciferase reporter assays. Insect cell expression of RBBP for crystallisation was carried out by CD in Sydney Analytical with the help of JPM. VTT and OV synthesized peptides and performed fluorescence polarization experiments. FB conducted lf-qMS experiments. TF and MB bioinformatically analysed next-generation sequencing data with the help of JiLa, LP and TMW. SBH and TMW wrote the manuscript with support and help from all other co-authors.

## Funding

This work was supported by the Deutsche Forschungsgemeinschaft (DFG) through the Collaborative Research Center TRR 81 (project A15) and the Cardio Pulmonary Institute (CPI) to SBH.

## Conflict of interest statement

None declared.

## Notes

### Competing Interest Statement

The authors have declared no competing interest.

