## Supplementary Information for "ZNF512B binds RBBP4 via a variant NuRD interaction motif and aggregates chromatin in a NuRD complex-independent manner"

[§]shared authors

[&]Current address: Institute of Biochemistry, Justus-Liebig University Giessen, 35392 Giessen, Germany

[*]Corresponding author: Sandra B. Hake, Institute for Genetics, Justus-Liebig-University Giessen, Heinrich-Buff-Ring 58-62, 35392 Giessen, Germany, EMAIL:, phone: 0049 (0)641 99 35460, FAX: 0049 (0)641 99 35469

Running title: ZNF512B is a NuRD binder and chromatin aggregator

Key words: ZNF512B / H2A.Z / RBBP4 / NuRD / chromatin compaction / zinc finger

### Supplementary Materials and Methods

#### *Antibodies*

| Antibody | Host | Supplier | Order Number | Application | Dilution |
|---|---|---|---|---|---|
| α-CHD4 | Mouse | Abcam | ab70469 | IF | 1:100 |
| α-FLAG | Mouse | Sigma-Aldrich | F3165 | WB | 1:6.000 |
| α-GFP | Mouse | Roche | 11814460001 | WB | 1:3.000 |
| α-H2A.Z | Rabbit | Abcam | ab4174 | WB | 1:1.000 |
| α-H2A.Zac | Rabbit | Abcam | ab232908 | IF | 1:100 |
| α-H3 | Rabbit | Abcam | ab1791 | WB | 1:5.000 |
| α-H3K27ac | Rabbit | Active Motif | 39133 | IF | 1:100 |
| α-H3K27me3 | Rabbit | Diagenode | C15410195 | IF | 1:100 |
| α-H3K4me3 | Rabbit | Diagenode | C15410003 | IF | 1:100 |
| α-H3K9me3 | Rabbit | Invitrogen | 49-1008 | IF | 1:100 |
| α-H3S10ph | Rabbit | Invitrogen | PA5-17869 | IF | 1:100 |
| α-HDAC1 | Rabbit | Proteintech | 10197-1-AP | WB | 1:1.000 |
| α-HDAC2 | Rabbit | Abcam | ab7029 | WB | 1:1.000 |
| α-HMG20A | Rabbit | Proteintech | 12085-1-AP | WB | 1:1.000 |
| α-KPNA4 | Rabbit | Proteintech | 12463-1-AP | WB | 1:1.000 |
| α-MBD2 | Rabbit | Abcam | ab188474 | WB | 1:1.000 |
| α-MTA1 | Rabbit | Cell Signaling Technology | 5647 | WB | 1:1.000 |
| | | | | IF | 1:100 |
| α-PWWP2A | Rabbit | Norvusbio | NBP2-13833 | WB | 1:1.000 |
| α-RBBP4 | Rabbit | Abcam | ab79416 | WB | 1:1.000 |
| | | | | IF | 1:100 |
| α-ZNF512B | Rabbit | BJ-Diagnostik BioScience GmbH | not applicable | WB | 1:1000 |
| | | | | IF | 1:100 |
| anti-Mouse IgG (H+L), HRP | Goat | Invitrogen | 31430 | WB | 1:20.000 |
| anti-Rabbit IgG (H+L), HRP | Goat | Invitrogen | 31460 | WB | 1:20.000 |
| F(ab')2 anti-Rabbit IgG (H+L) Cross-Adsorbed, Alexa Fluor™ 488 | Goat | Invitrogen | A-11070 | IF | 1:200 |
| F(ab')2 anti-Rabbit IgG (H+L) Cross-Adsorbed, Alexa Fluor™ 594 | Goat | Invitrogen | A-11072 | IF | 1:200 |

| | | | | | |
|---|---|---|---|---|---|
| F(ab')2 anti-Mouse IgG (H+L) Cross-Adsorbed, Alexa Fluor™ 594 | Goat | Invitrogen | A-11020 | IF | 1:200 |

#### *Primers and Oligos*

| Target | Application | Forward (5' to 3') | Reverse (5' to 3') |
|---|---|---|---|
| ZNF512B | Cloning into pIRESneo-eGFP | TCATCGTTCGAAGTCCGGATCTATGACGGATCCTTTCTGCGTTGGAG | TAGTAAGCGGCCGCTCACTATCACTTTTCAGGCGCCTTGCTG |
| | Cloning into pEGFP-N2 | ATGACGGATCCTTTCTGCGTTGGAG | CTTTTCAGGCGCCTTGCTGACTC |
| | Cloning into p3xFLAG-CMV-10 | ATGACGGATCCTTTCTGCGTTGGAG | TCACTTTTCAGGCGCCTTGCTG |
| | Cloning into pFastBac1 | ACTACTGAATTCATGGATTACAAGGATGACGATGACAAGGGTGGTTCTGGTACGGATCCTTTCTGCGTTGGAG | GTGGTGGGTACCTCACTTTTCAGGCGCCTTGCT |
| | Cloning into pAB-Gal94 | TTTGTCGACACGGATCCTTTCTGCGTTGGA | TTTTTCTAGACTACTTTTCAGGCGCCTTGCTGA |
| ZNF512B | Cloning of GFP-ZNF512B_ΔZF | TCATCGTTCGAAGTCCGGATCTATCAGCAGGCCGGTCACCATC | TAGTAAGCGGCCGCTCACTACTCTTCAGGGCCACCTGGAG |
| | Cloning of GFP-ZNF512B_ΔI fragment 1 | TCATCGTTCGAAGTCCGGATCTATGACGGATCCTTTCTGCGTTGGAG | CACCTGGAGCGATGGTGACCGGCCTGCT |
| | Cloning of GFP-ZNF512B_ΔI fragment 2 | GGTCACCATCGCTCCAGGTGGCCCTGAA | TAGTAAGCGGCCGCTCACTACACCTTCCGCTTCTTCTTGGGCTTTTCAGGCGCCTTGCTGACTC |
| | Cloning of GFP-ZNF512B_ΔZF1-2 | TCATCGTTCGAAGTCCGGATCTATCAGCAGGCCGGTCACCATC | TAGTAAGCGGCCGCTCACTATCACTTTTCAGGCGCCTTGCTG |
| | Cloning of GFP-ZNF512B_ΔZF3-8 | TCATCGTTCGAAGTCCGGATCTATGACGGATCCTTTCTGCGTTGGAG | TAGTAAGCGGCCGCTCACTACTCTTCAGGGCCACCTGGAG |
| ZNF512B | Cloning of GAL-ZNF512B_ΔZF Gibson vector | CAGGTGGCCCTGAAGAGTGATTTCGGATCCAAAGCTTGATCCG | ATGGTGACCGGCCTGCTGATGAATTCCAATCTAGATTGCGGCG |
| | Cloning of GAL-ZNF512B_ΔZF Gibson fragment | CGCAATCTAGATTGGAATTCATCAGCAGGCCGGTCAC | ATCAAGCTTTGGATCCGAAATCACTCTTCAGGGCCACCT |
| | Cloning of GAL-ZNF512B_ΔI Gibson vector | GAGTCAGCAAGGCGCCTGAATTTCGGATCCAAAGCTTGATCG | CCAACGCAGAAAGGATCCGTGAATTCCAATCTAGATTGCGGC |
| | Cloning of GAL-ZNF512B_ΔI Gibson fragment | CGCAATCTAGATTGGAATTCACGGATCCTTTCTGCGTTGG | ATCAAGCTTTGGATCCGAAATTCAGGCGCCTTGCTGAC |
| ZNF512B | SDM K419A | GCGCACAGAAGGAAACAGAAAACACCCAAAAAGTTTACAGGGGAGC | TTTCTGTTTCCTTCTGTGCGCTGTGCGCTCCGGGTC |
| | SDM H420A | AGGCCAGAAGGAAACAGAAAACACCCAAAAAGTTTACAGGGGAGC | TTTCTGTTTCCTTCTGGCCTTTGTGCGCTCCGGGTC |
| | SDM R421A | AGCACGCAAGGAAACAGAAAACACCCAAAAAGTTTACAGGGGAGC | TTTCTGTTTCCTTGCGTGCTTTGTGCGCTCCGGGTC |
| | SDM R422A | AGCACAGAGCGAAACAGAAAACACCCAAAAAGTTTACAGGGGAGC | TTTCTGTTTCGCTCTGTGCTTTGTGCGCTCCGGGTC |

| | | | |
|---|---|---|---|
| | SDM K423A | CACAGAAGGGCACAGAAAACACCCAAAAAGTTTACAGGGGAGCAGC | GTTTTCTGTGCCCTTCTGTGCTTTGTGCGCTCCGGGTCCT |
| | SDM Q424A | AGCACAGAAGGAAAGCGAAAACACCCAAAAAGTTTACAGGGGAGC | TTTCGCTTTCCTTCTGTGCTTTGTGCGCTCCGGGTC |
| | SDM K425A | AGCACAGAAGGAAACAGGCAACACCCAAAAAGTTTACAGGGGAGC | TGCCTGTTTCCTTCTGTGCTTTGTGCGCTCCGGGTC |
| | SDM T426A | AGCACAGAAGGAAACAGAAAGCACCCAAAAAGTTTACAGGGGAGC | TTTCTGTTTCCTTCTGTGCTTTGTGCGCTCCGGGTC |
| | SDM P427A | AGCACAGAAGGAAACAGAAAACAGCCAAAAAGTTTACAGGGGAGC | TTTCTGTTTCCTTCTGTGCTTTGTGCGCTCCGGGTC |
| | SDM K428A | AGCACAGAAGGAAACAGAAAACACCCGCAAAGTTTACAGGGGAGC | TTTCTGTTTCCTTCTGTGCTTTGTGCGCTCCGGGTC |
| | SDM K429A | AGCACAGAAGGAAACAGAAAACACCCAAAGCGTTTACAGGGGAGC | TTTCTGTTTCCTTCTGTGCTTTGTGCGCTCCGGGTC |
| | SDM K429R | AGCACAGAAGGAAACAGAAAACACCCAAAAGGTTTACAGGGGAGC | TTTCTGTTTCCTTCTGTGCTTTGTGCGCTCCGGGTC |
| | SDM F430A | AGCACAGAAGGAAACAGAAAACACCCAAAAAGGCTACAGGGGAGC | TTTCTGTTTCCTTCTGTGCTTTGTGCGCTCCGGGTC |
| | SDM K419A_R421A | GCGCACGCAAGGAAACAGAAAACACCCAAAAAGTTTACAGGGGAGCAG | GTTTCCTTGCGTGCGCTGTGCGCTCCGGGTCC |
| ZNF512B | RT-qPCR | TCCCAACGACTGCTGTGAAG | TGAACTCCTTCGGACACAGC |
| H19 | EMSA | CACCCGGTGCTTCGGGCCCTCTAGCCCGGGCTTTTTCTAACTGGAGTGGCTCCGCCCA | |

***RBBP4-ZNF512B-Structure: Data collection and refinement statistics***

| **Data collection** | |
|---|---|
| Space group | P 1 21 1 |
| Cell dimensions<br>a, b, c (Å)<br>α, β, γ (°) | 76.10, 59.58, 101.45<br>90, 93.77, 90 |
| Resolution (Å) | 46.88 - 2.20 (2.32 - 2.20) |
| $R_{merge}$ | 0.156 (1.238) |
| $CC_{1/2}$ | 0.991 (0.493) |
| $I / \sigma I$ | 6.5 (1.3) |
| Completeness (%) | 100 (100) |
| Redundancy | 3.5 (3.6) |
| **Refinement** | |
| Resolution (Å) | 46.88 - 2.20 |
| No. reflections | 46302 (4563) |
| $R_{work}$/ $R_{free}$ | 0.21 (0.33)/ 0.24 (0.35) |
| Ramachandran statistics<br>Favoured (%)<br>Allowed (%)<br>Outliers (%) | 97.11<br>2.63<br>0.26 |
| Number of non-hydrogen atoms<br>Macromolecules<br>Ligands<br>Solvent | 6185<br>161<br>337 |
| Average *B*-factor | 41.40 |
| R.m.s deviations | |
| Bond lengths (Å) | 0.005 |
| Bond angles (°) | 0.80 |

### Supplementary Figures S1-S6 and Legends

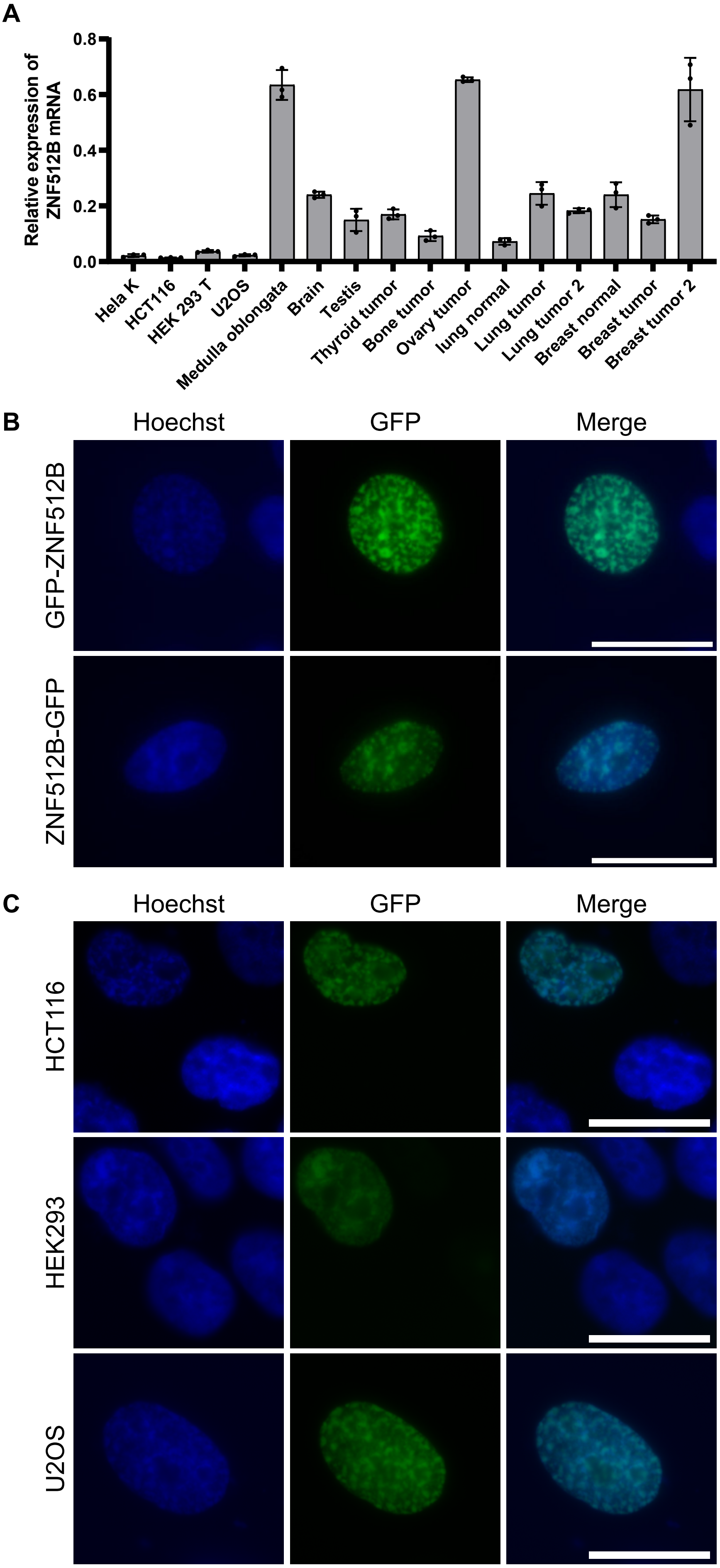
A
Relative expression of ZNF512B mRNA
0.8
0.6
0.4
0.2
0.0
Hela K
HCT116
HEK 293 T
U2OS
Medulla oblongata
Brain
Testis
Thyroid tumor
Bone tumor
Ovary tumor
lung normal
Lung tumor
Lung tumor 2
Breast normal
Breast tumor
Breast tumor 2
B
Hoechst
GFP
Merge
GFP-ZNF512B
ZNF512B-GFP
C
Hoechst
GFP
Merge
HCT116
HEK293
U2OS

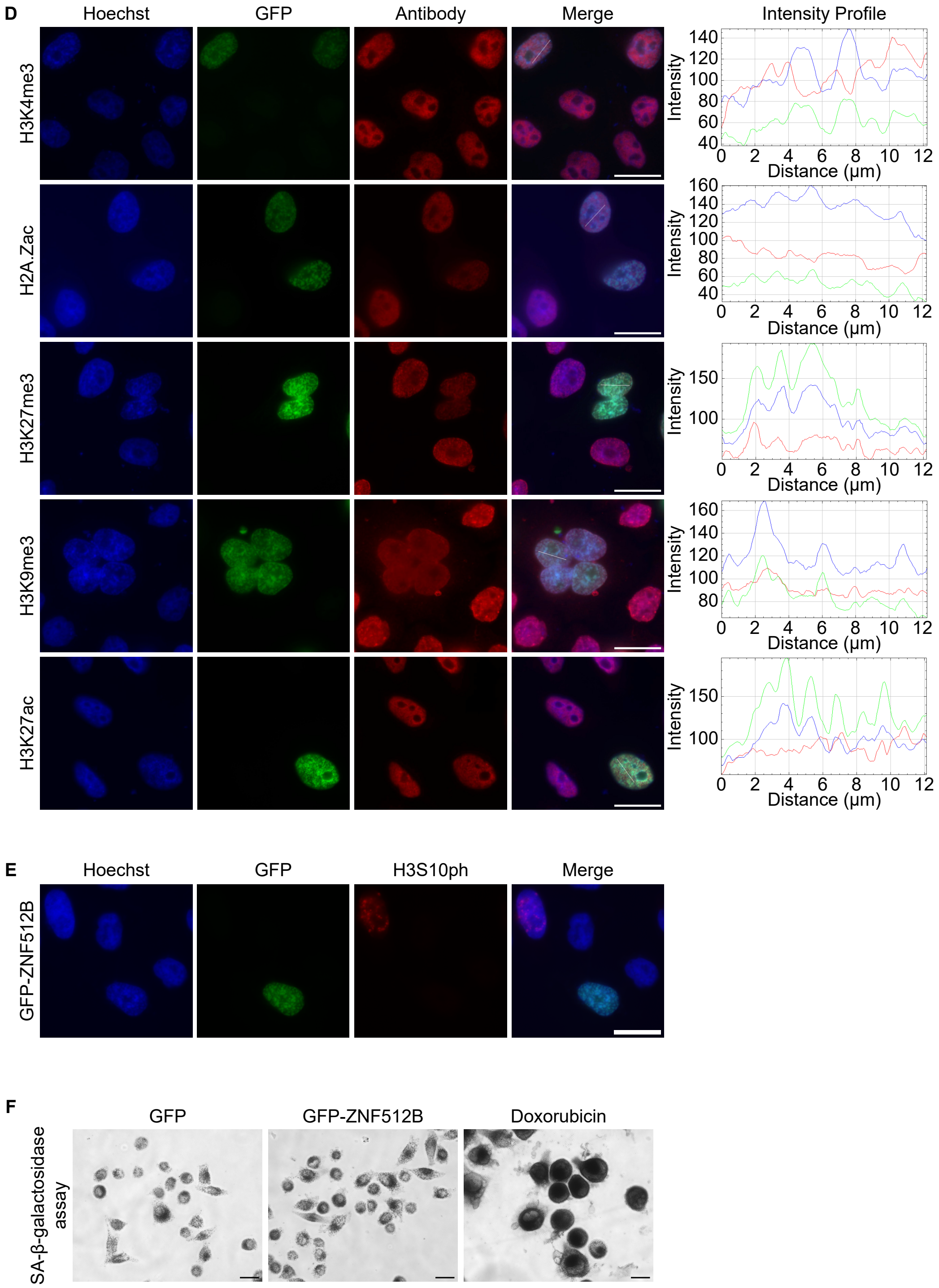

**Supplementary Figure S1: Chromatin aggregation due to ZNF512B's overexpression is independent of tag localization, cell type and cell cycle. (A)** RT-qPCR analysis of ZNF512B mRNA expression in different human cell lines and tissues normalized to HPRT1 expression. SD = three technical replicates. **(B-C)** Immunofluorescence microscopy of Hoechst (DNA, blue) stained **(B)** HeLaK cells expressing GFP-ZNF512B or ZNF512B-GFP (green), **(C)** U2OS, HCT116 or HEK239 cells expressing GFP-ZNF512B (green). Scale bars: 20 µm. **(D, E)** Left: IF microscopy of Hoechst (DNA, blue) stained HeLaK cells expressing GFP-ZNF512B (green) and co-stained with **(D)** different antibodies against active (H3K4me3, H3K27ac, H2A.Zac) or repressive (H3K9me3, H3K27me3) histone modifications (red) or **(E)** anti-H3S10ph antibody (red) as mitosis mark. Scale bars: 20 µm. Right: Intensity profile of nuclear areas (see lines, left) depicting DNA (blue), GFP-ZNF512B (green) and respective histone PTMs (red) fluorescence. **(F)** Bright-field microscopy of *in situ* staining for β-galactosidase activity in HeLaK cells expressing GFP or GFP-ZNF512B as marker for senescent cells. Pre-treatment of HeLaK cells with Doxorubicin served as positive control. Scale bar = 20 µm.

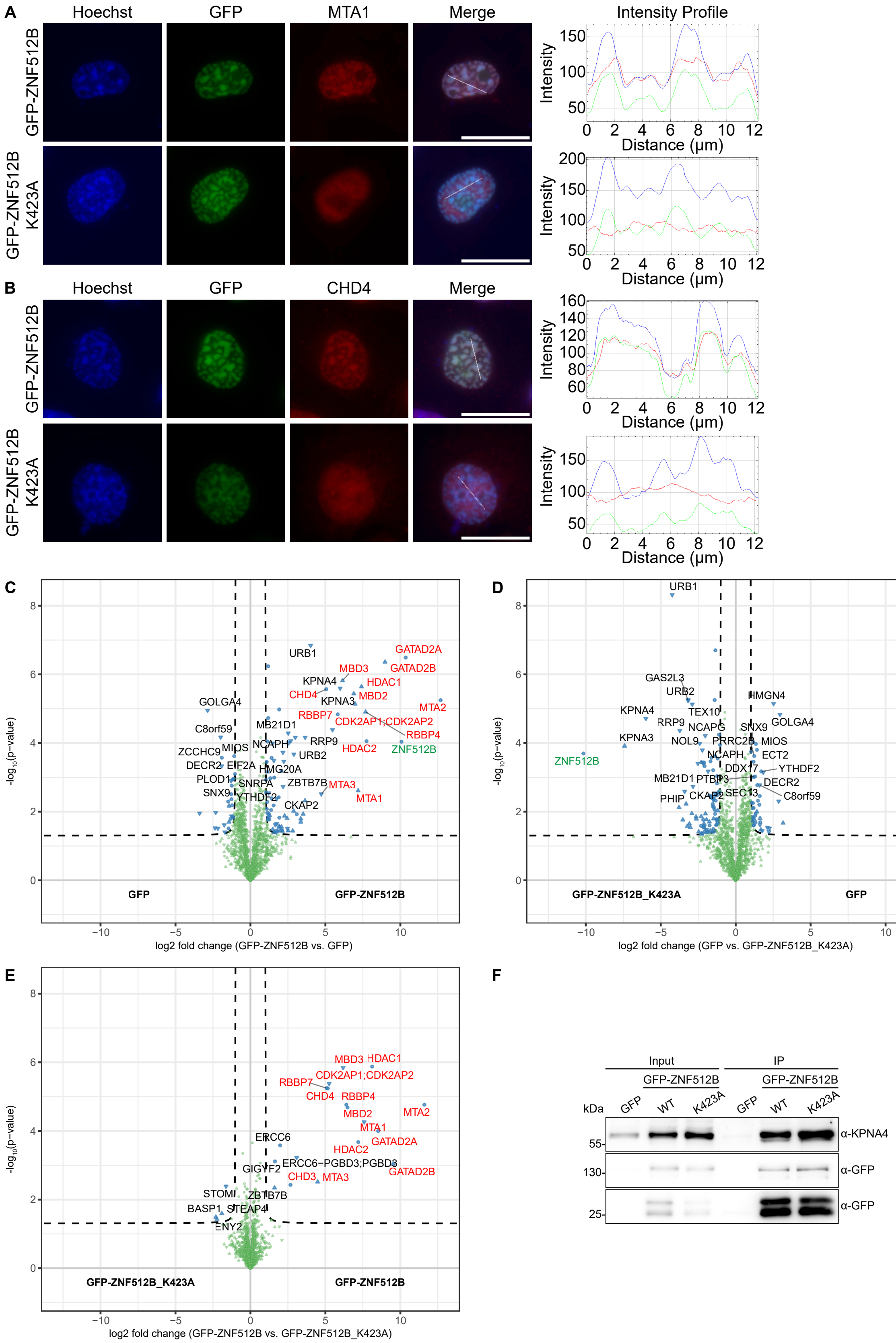

**Supplementary Figure S2: ZNF512B interaction with NuRD depends on its NIM.** **(A, B)** Immunofluorescence microscopy of HeLaK cells expressing GFP or GFP-ZNF512B_K423A (green) co-stained with Hoechst (DNA, blue) and antibody against

MTA1 **(A)** or CHD4 **(B)** (red). Scale bar = 20 µm. **(C-E)** Volcano plots of lf-qMS data (one replicate) comparing proteins enriched on GFP (control) with those bound to GFP-ZNF512B **(C),** or those bound to GFP-ZNF512B_K423A with GFP **(D)** or those bound to GFP-ZNF512B_K423A with GFP-ZNF512B **(E).** ZNF512B protein is highlighted in green, NuRD members in red and other binding proteins in black. See also Figure 2D for heatmap. **(F)** Immunoblots of nuclear extracts from HeLaK cells transiently expressing GFP, GFP-ZNF512B or GFP-ZNF512B_K423A after pull down with GFP-TRAP beads detecting NIM-independent binding of KPNA4.

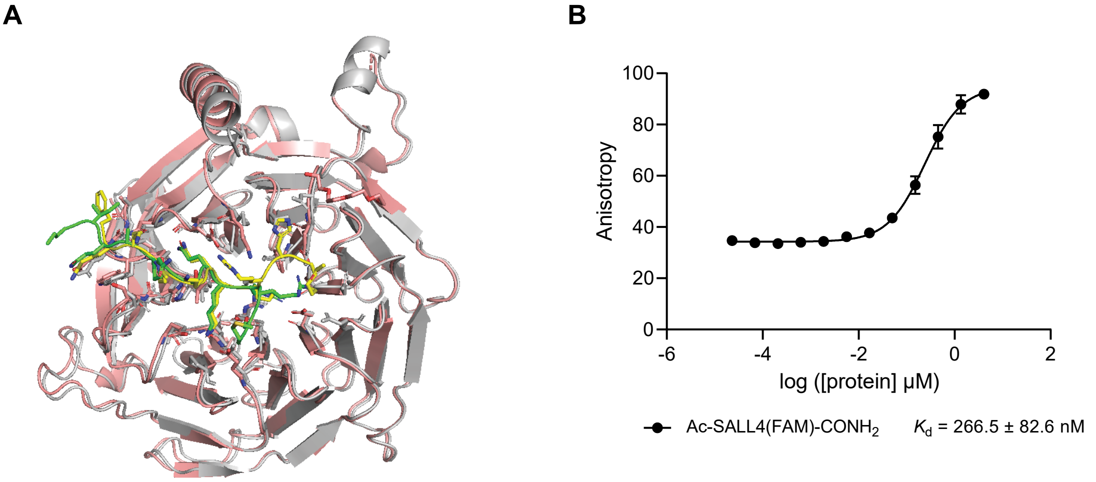

**Supplementary Figure S3**. **Binding of the ZNF512B NIM to RBBP4. (A)** Overlay (over backbone heavy atoms) of the RBBP4-ZNF512B structure (grey and yellow) with the RBBP4-FOG-1 structure (salmon and green, PDB: 2XU7, (1)). **(B)** Fluorescence polarization-based binding experiments of RBBP4. Shown is a saturation binding experiment of 1 nM fluorescently labelled Ac-SALL4(2-12)-FAM in 50 mM Tris-Cl pH 7.5, 150 mM NaCl, 0.02% Triton X-100.

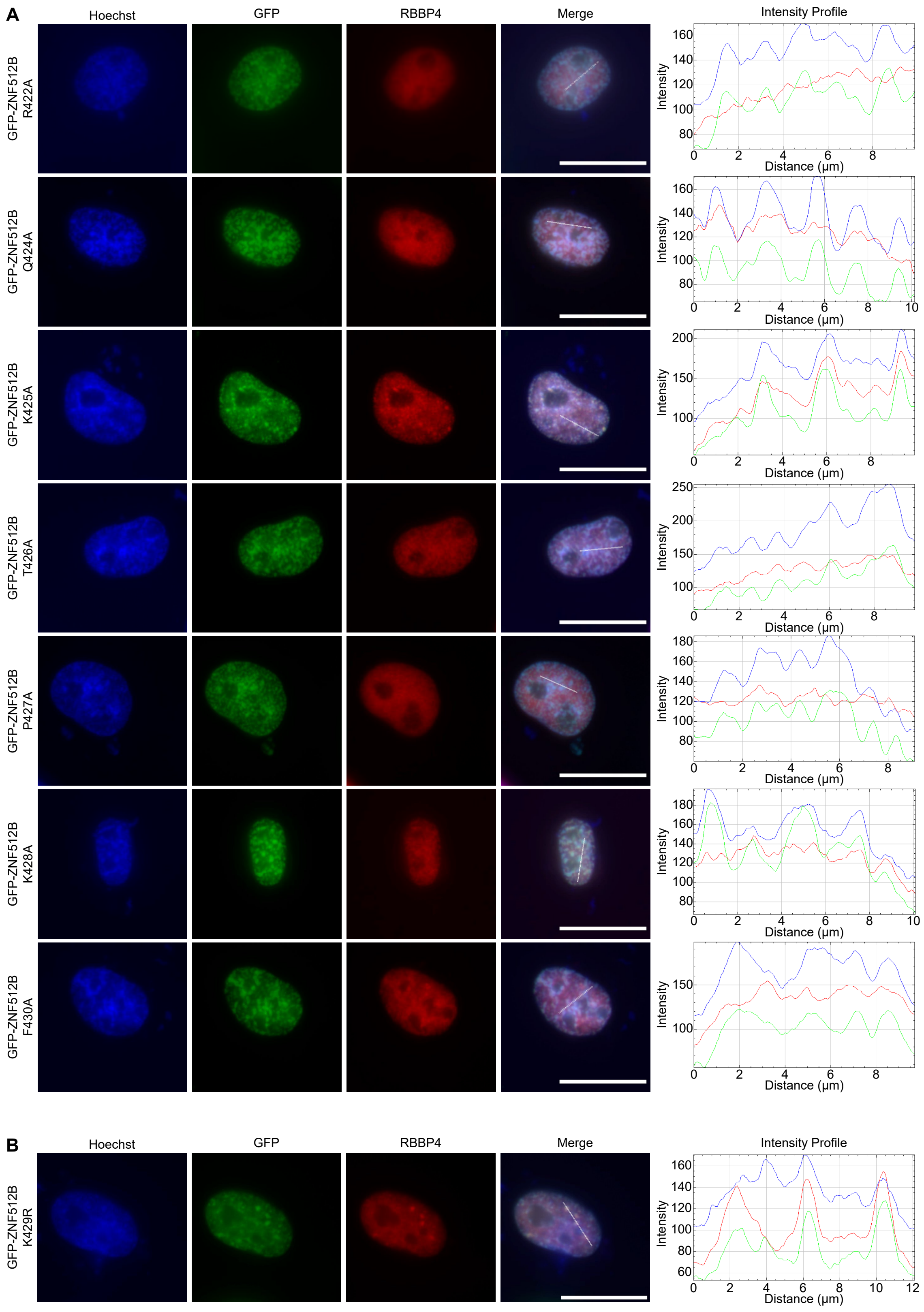

**Supplementary Figure S4: ZNF512B contains a functional internal NIM. (A, B)** Left: Immunofluorescence microscopy of HeLaK cells expressing **(A)** GFP-ZNF512B NIM alanine mutants or **(B)** GFP-ZNF512B_K429R co-stained with Hoechst (DNA, blue) and anti-RBBP antibody (red). Scale bar: 20 µm. Right: Intensity profile of nuclear areas (see lines, left) depicting DNA (blue), GFP-ZNF512B mutants (green) and RBBP4 (red) fluorescence.

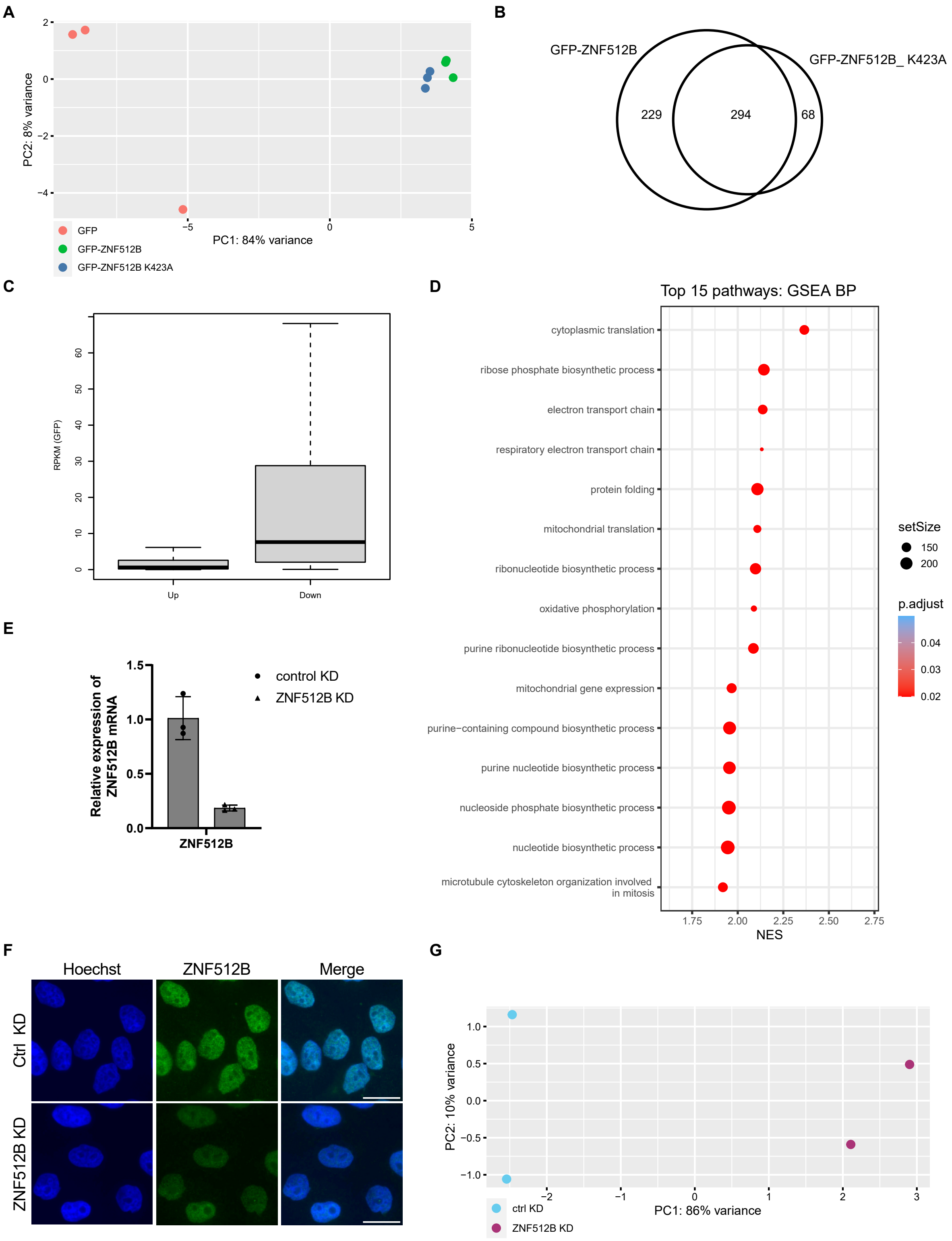

A
PC2: 8% variance
PC1: 84% variance
GFP
GFP-ZNF512B
GFP-ZNF512B K423A
B
GFP-ZNF512B
GFP-ZNF512B_ K423A
229
294
68
C
RPKM (GFP)
Up
Down
D
Top 15 pathways: GSEA BP
cytoplasmic translation
ribose phosphate biosynthetic process
electron transport chain
respiratory electron transport chain
protein folding
mitochondrial translation
ribonucleotide biosynthetic process
oxidative phosphorylation
purine ribonucleotide biosynthetic process
mitochondrial gene expression
purine−containing compound biosynthetic process
purine nucleotide biosynthetic process
nucleoside phosphate biosynthetic process
nucleotide biosynthetic process
microtubule cytoskeleton organization involved in mitosis
NES
setSize
150
200
p.adjust
0.04
0.03
0.02
E
Relative expression of ZNF512B mRNA
control KD
ZNF512B KD
ZNF512B
F
Hoechst
ZNF512B
Merge
Ctrl KD
ZNF512B KD
G
PC2: 10% variance
PC1: 86% variance
ctrl KD
ZNF512B KD

H

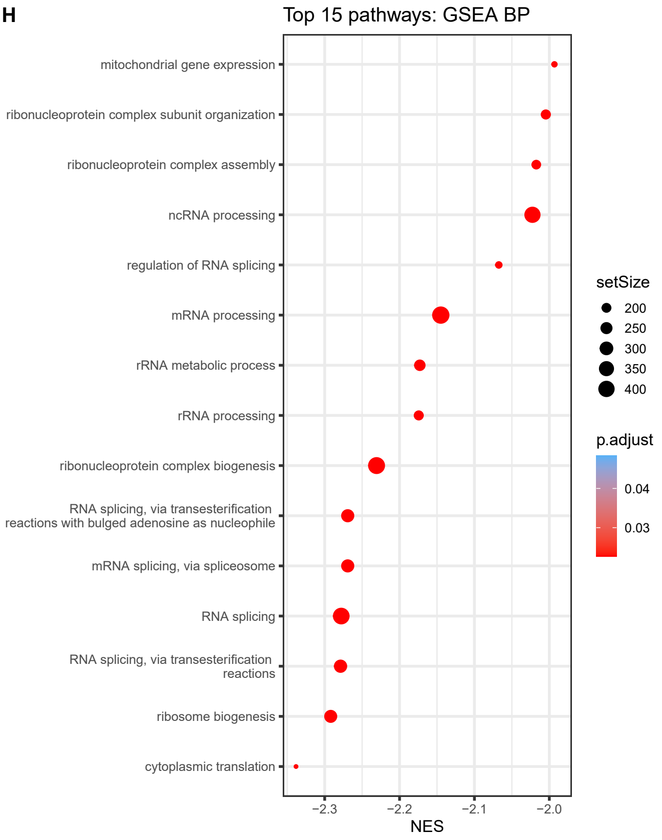

I

**BIRC3**
**CDC14A**
**CREB5**
**IL6**
**NFKBIE**
**PTGS2**
**RELB**
**TMEM52B**
**TNFAIP3**

**Supplementary Figure S5: Gene expression changes upon GFP-ZNF512B or GFP-ZNF512B_K423A overexpression or ZNF512B depletion. (A)** Principal component analysis (PCA) of RNA-seq data of three replicates of GFP (red), GFP-ZNF512B (green) or GFP-ZNF512B_K423A (blue) expressing HeLaK cells. **(B)** Euler diagram depicting the overlap of significantly deregulated genes (log2 FC > 1 or < -1 and adjusted p-value < 0.05) upon GFP-ZNF512B or GFP-ZNF512B_K523A overexpression compared to GFP control. **(C)** Box plot depicting gene expression base levels (RPKM) of HeLaK cells upon GFP-ZNF512B or GFP-ZNF512B_K423A overexpression. Shown are genes significantly deregulated divided in upregulation or downregulation **(D)** Gene set enrichment analysis (GSEA) for the Gene Ontology (GO) database of deregulated genes upon GFP-ZNF512B and GFP-ZNF512B_K423A overexpression. **(E)** RT-qPCR to detect relative ZNF512B mRNA expression upon control (Ctrl) or ZNF512B siRNA-mediated knock-down (KD) in HeLaK cells normalized to HPRT1 expression. SD = three biological replicates. **(F)** Immunofluorescence microscopy of HeLaK cells upon control (Ctrl) or ZNF512B siRNA-mediated knock-down (KD) co-stained with anti-ZNF512B antibody (green) and Hoechst (DNA, blue). Scale bar = 20 µm. **(G)** Principal component analysis (PCA) of RNA-seq data of two replicates of control (Ctrl) or ZNF512B siRNA-mediated KD

HeLaK cells. **(H)** Gene set enrichment analysis (GSEA) for the Gene Ontology (GO) database of deregulated genes upon ZNF512B knock-down (KD). **(I)** List of genes downregulated upon GFP-ZNF512B overexpression and upregulated upon ZNF512B knockdown.

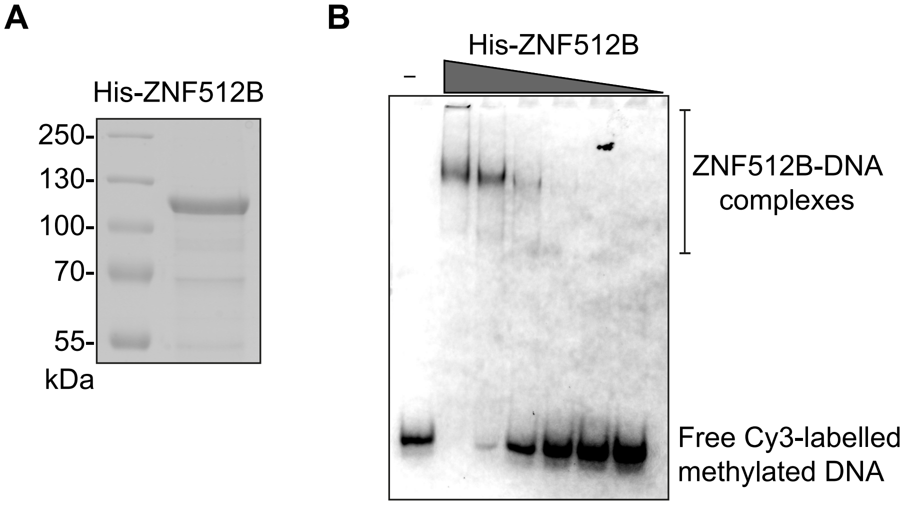

**Supplementary Figure S6: ZNF512B overexpression does not affect global chromatin compaction. (A)** Coomassie brilliant blue-stained SDS-PAGE gel separating purified recombinant His-ZNF512B. **(B)** Electrophoretic Mobility Shift Assay (EMSA) of Cy3-labelled methylated DNA together with increasing amounts of purified recombinant His-ZNF512B protein**.**
